# Trans-allelic Epigenetic Dominance Disrupts Hybrid Endosperm Development in Wild Tomatoes

**DOI:** 10.64898/2026.03.15.711876

**Authors:** Ana Marcela Florez-Rueda, Morgane Roth, Thomas Städler

## Abstract

**Background and Aims:** Hybrid seed failure (HSF) is a common reproductive barrier in flowering plants, but how divergence in so-called ‘effective ploidy’ translates into genome-wide allelic imbalance in hybrid endosperm remains unresolved. Using wild tomatoes (*Solanum* sect. *Lycopersicon*) as our model system, we test whether parent-of-origin expression shifts in hybrids are consistent with simple dosage effects or instead reflect lineage-specific *trans* regulation.

**Methods:** Using reciprocal interspecific crosses, we quantified parent-of-origin (allele-specific) expression in developing endosperm and analyzed patterns of differential parent-of-origin expression (DPE) across cross contexts. Guided by strong patterns in parental expression, we further classified DPE genes into four functional classes defined by their relationship to *Solanum peruvianum* (*Per*). This exercise captured distinct modes of *trans*-allelic regulation and guided functional interpretation.

**Key Results:** Parental expression proportions are strongly asymmetric in hybrids, with the most pronounced and repeatable shifts occurring in crosses involving *Per*. These shifts recur across reciprocal contexts and include elevated maternal proportions even in crosses phenotypically classified as paternal-excess-like, arguing against a dosage-only model. Instead, we observed cross-consistent, coordinated DPE patterns corresponding to *trans*-acting dominance associated with *Per*. Functional enrichment of *Per*-associated DPE highlights chromatin regulation, including DNA methylation/RdDM- and chromatin remodeling-related factors, and Polycomb-linked regulators, implicating disruption of chromatin-based repression in hybrid endosperm. Concurrently, *Per*-associated activation of auxin- and cell-cycle regulatory pathways in non-*Per* alleles suggests mis-timed hormone-dependent developmental transitions that can destabilize proliferation–cellularization programs.

**Conclusions:** Our findings support a model of *trans*-allelic epigenetic dominance in which *Per*-associated regulatory inputs reshape allelic expression landscapes in hybrid endosperm largely independent of parental origin. This provides a mechanistic link between effective ploidy divergence, genome-wide transcriptional imbalance, and HSF, and motivates reciprocal hybrid studies that integrate expression with chromatin-state and accessibility profiling.

## INTRODUCTION

The evolutionary success of angiosperms relies on the coordinated development of reproductive tissues within the flower and, ultimately, the seed, that differ in both ploidy and parental genomic contributions. Following double fertilization, the embryo and endosperm that constitute the developing seed, inherit the same parental genomes but in distinct ratios, with the endosperm typically exhibiting a maternal genomic excess (2m:1p). This unique genomic configuration renders endosperm development particularly sensitive to gene dosage and parental balance, necessitating fine-tuned regulatory mechanisms to ensure proper growth and resource allocation (Baroux et al. 2002).

Hybrid seed failure (HSF) is widely understood as a consequence of disrupted parental genome balance in the endosperm rather than malfunction of a single epigenetic pathway. Classical and contemporary models rooted in parental-conflict theory posit that divergent evolutionary pressures on maternally and paternally inherited genomes generate dosage-sensitive regulatory incompatibilities, particularly in the endosperm, a tissue whose development critically depends on precise genomic ratios and parent-of-origin–specific gene expression (Haig and Westoby 1989, 1991; Brandvain and Haig 2005; Lafon-Placette and Köhler 2016; Povilus et al. 2018; Coughlan et al. 2020; Köhler et al. 2021; Sandstedt and Sweigart 2022). Consistent with this view, developing seeds from interspecific and interploidy crosses frequently exhibit misregulation of imprinted genes, many of which are dosage sensitive and controlled by DNA methylation and histone-modification asymmetries that are established in the gametes (Gehring and Satyaki 2017; Batista and Köhler 2020). Perturbation of these epigenetic marks can lead to widespread deregulation of imprinted loci and, critically, to secondary trans-acting effects on broader transcriptional networks that extend beyond imprinted genes themselves, amplifying developmental imbalance and precipitating endosperm failure (Florez-Rueda et al. 2016; Kirkbride et al. 2019; Satyaki and Gehring 2019). Thus, HSF is best viewed as an emergent property of disrupted parental dosage control and epigenetic regulation in the endosperm, with multiple layers of chromatin-based regulation contributing to hybrid incompatibility (Baroux et al. 2007; Martienssen 2010; Dziasek et al. 2021; Minow et al. 2025).

A central conceptual framework for understanding HSF is the sensitivity of endosperm development to parental genome dosage and balance (Haig and Westoby 1989). Classical work on interploidy crosses established the concepts of maternal excess (ME) and paternal excess (PE), in which deviations from the canonical 2 maternal:1 paternal genome ratio in the endosperm lead to predictable and reproducible developmental defects, including premature cellularization under ME and prolonged nuclear proliferation under PE. These phenotypes are particularly well characterized in *Arabidopsis thaliana*, where reciprocal diploid–tetraploid crosses (2x × 4x) generate strong, direction-dependent endosperm failure and transcriptional misregulation, underscoring the dosage sensitivity of parent-of-origin–dependent regulatory networks (Scott et al. 1998; Köhler et al. 2010; Lafon-Placette and Köhler 2015). Importantly, analogous outcomes are also observed in homoploid interspecific crosses in *Arabidopsis* and *Solanum*, where parents share the same chromosome number but nevertheless exhibit ME-like and PE-like phenotypes, indicating that functional dosage differences can arise independently of karyotypic ploidy (Lafon-Placette et al. 2017; Roth et al. 2018a). This insight is formalized in the concept of ‘effective ploidy’, which captures lineage-specific differences in the regulatory contribution of parental genomes to endosperm development (Lafon-Placette and Köhler 2016; Städler et al. 2021). The fully equivalent but earlier framework ‘endosperm balance number’ (EBN), developed from studies of wild potatoes (*Solanum*), demonstrated that successful seed development depends on achieving a balanced maternal:paternal EBN ratio rather than matching physical ploidy *per se*, thereby explaining why certain homoploid crosses fail while others succeed (Johnston et al. 1980). These concepts establish parental dosage imbalance—whether generated by interploidy crosses or divergence in effective ploidy—as a unifying mechanism underlying hybrid endosperm failure.

Among the molecular pathways implicated in HSF, epigenetic mechanisms that regulate parental genome expression in the endosperm have received particular attention. The RNA-directed DNA methylation (RdDM) pathway has emerged as an important modulator of parental dosage sensitivity, with genetic and transcriptomic analyses in *Arabidopsis* and *Brassica* demonstrating that perturbation of canonical RdDM components—especially paternally acting factors—can strongly influence hybrid seed viability and endosperm gene expression (Grover et al. 2018; Kirkbride et al. 2019; Satyaki and Gehring 2019). However, recent molecular evidence indicates that epigenetic control of plant development extends beyond RdDM, encompassing higher-order chromatin organization and *cis*-regulatory architecture as key determinants of transcriptional regulation (Marand et al. 2023; Yan et al. 2024; Xu et al. 2025). Distinct chromatin organization features between endosperm and vegetative tissues have been documented in *Arabidopsis*, where long-range chromatin interactions and looping correlate with transcriptional activity and the regulation of imprinted loci, highlighting a role for three-dimensional genome architecture in shaping tissue-specific expression programs relevant to seed development and viability (Yadav et al. 2021). In parallel, chromatin accessibility profiling has revealed extensive accessible regions in plant genomes, including intergenic *cis*-regulatory elements enriched for transcription-factor-binding motifs, underscoring the importance of accessible chromatin in recruiting regulatory proteins and coordinating developmental transcriptional networks (Marand et al. 2023; Yan et al. 2024). Consistent with this multilayered regulatory framework, dynamic histone modification landscapes—comprising both activating (e.g. H3K4me3, H3K9ac) and repressive marks (e.g. H3K27me3)—are tightly associated with gene expression changes during seed and endosperm development across plant species, indicating that chromatin-state transitions and remodeling contribute broadly to transcriptional control in reproductive tissues (He et al. 2024).

Due to their pronounced differences in effective-ploidy and well-characterized hybrid endosperm phenotypes, wild tomatoes (*Solanum* section *Lycopersicon*) provide exceptional opportunities to investigate these questions (Roth et al. 2018a). Previous work in *Solanum* has revealed extensive transcriptional dysregulation in hybrid endosperm, with predominantly reduced gene expression relative to intraspecific crosses and asymmetric expression patterns depending on cross direction (Florez-Rueda 2014; Florez-Rueda et al. 2016; Roth et al. 2019). Notably, Maternally Expressed Genes (MEGs) are preferentially overexpressed in ME-like crosses, whereas Paternally Expressed Genes (PEGs) are overexpressed in PE-like crosses, consistent with quantitative effects proportional to effective-ploidy divergence (Roth et al. 2018b, 2019; Städler et al. 2021). In this system, *Solanum peruvianum* (*Per*) exhibits higher effective ploidy relative to *S. chilense* (*Chi*) and accessions from the Arcanum complex used here (*S. arcanum* var. *marañón* (*Ama*)), such that all crosses in which *Per* is the ovule parent yield ME-like hybrid endosperms, while the reciprocal crosses represent PE-like conditions (Roth et al. 2018a). Consistent with this asymmetry, *Per* has previously been shown to exert dominance in both gene expression and small-RNA abundance in hybrid endosperm (Florez-Rueda et al. 2021b).

Building on these studies, the present work integrates allele-specific expression analyses across a series of reciprocal interspecific hybrids with functional pathway interrogation to examine how effective-ploidy differences mediate parent-of-origin regulatory effects in hybrid endosperm. As summarized schematically in **Figure 1**, our experimental design leverages reciprocal crosses to disentangle parental role from genomic identity, allowing us to track how specific parental genomes exert asymmetric regulatory influences across hybrid contexts—a level of factual complexity that appears to not have been addressed in previous literature. By systematically comparing parent-specific allelic expression across multiple species-pair crosses, we identify a conserved set of genes whose expression is consistently altered in association with the *Per* genome, independent of its parental role. These patterns reveal distinct classes of asymmetric regulatory interactions between parental genomes, notably a mode of *trans*-allelic epigenetic dominance whereby the presence of *Per* reshapes hybrid regulatory landscapes of the opposite allele. By screening reciprocal crosses, we also identified widespread, systematic *cis* effects in which *Per* alleles themselves are activated or repressed. Functional analyses of *Per*-associated gene sets uncovered coordinated perturbations of epigenetic silencing machinery, histone-variant expression, and hormone-related pathways, linking *trans*-acting paternal dominance to disrupted chromatin regulation and transcriptional stability. Together, these findings establish *trans*-allelic epigenetic dominance of the higher-EBN lineage (here, *Per*) over lower-EBN lineages (here, *Ama* and *Chi*) as a unifying mechanism shaping hybrid endosperm dysfunction, and thus contributing to HSF.

**Figure 1.**
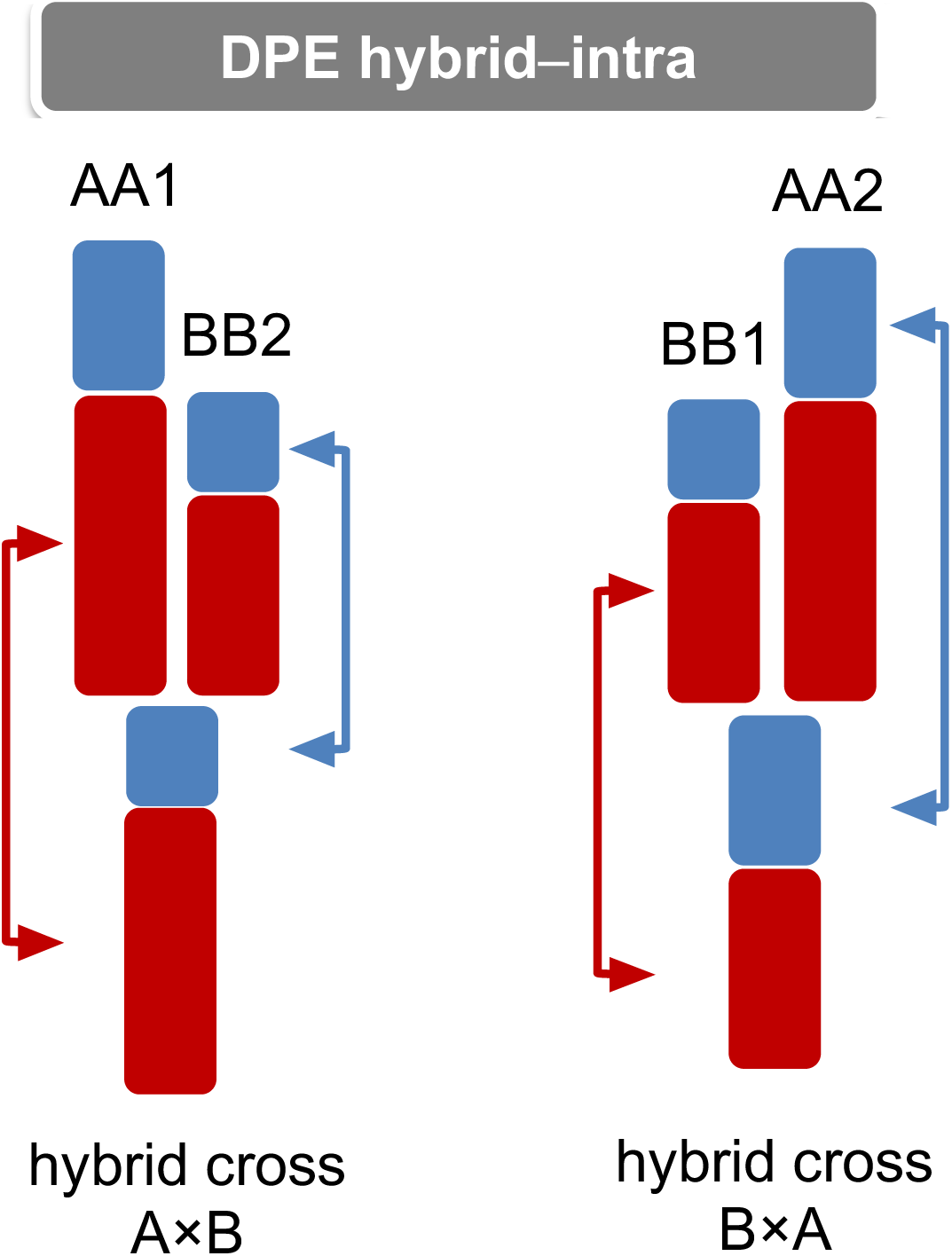
Schematic representation of formal expression comparisons in hybrid endosperms obtained from reciprocal crosses between species A and B (A×B and B×A). Differential Parental Expression (DPE) analyses hybrid–intra rest on comparisons between hybrid endosperms (at bottom of scheme) and each of their maternal and paternal references (i.e. normal within-species expression data, AA1, AA2, BB1, BB2). Heights of the colored bars reflect expression levels for a given gene and colors convey parent-specific expression (red for maternal, blue for paternal). Colored arrows symbolize the comparisons of parental expression proportions implemented in our study.

## RESULTS

Parent-specific expression in hybrid endosperms could be assessed for 11,195 to 11,985 genes (**Table S1**, **Dataset S1**), 9,728 of which could be evaluated in all three reciprocal hybrid crosses. The within-species data on parental expression proportions rest on 7,730 (lineage *Ama*), 9,623 (lineage *Chi*) and 13,198 genes (lineage *Per*; **Table S1**), reflecting the different levels of nucleotide polymorphism within the three species, as exemplified by the genome-wide differences of the chosen parental genotypes. Importantly, our tests of potential contamination from non-endosperm seed tissues (following Schon and Nodine 2017) reveal no evidence for such contamination in our laser-captured endosperm samples (**Figure S1**).

### Parental ratios in the endosperm of intra- and interspecific crosses

For all crosses and cross types, we first assessed parental contributions at genome-wide scales. We thus analyzed the distribution of per-gene maternal proportion, defined as the ratio between maternal and total expression (ranging from 0 to 1). Our data reveal marked differences in median maternal proportion between reciprocal intraspecific crosses in lineages *Ama* and *Chi*, while maternal proportions are almost identical for the two cross directions in lineage *Per* (**Table S1**). The expression data for the six hybrid crosses reveal that median maternal proportions vary with cross type and cross direction; the highest maternal proportions characterize the ME-like endosperms P×A and P×C, with medians of 0.809 and 0.801, respectively (**Figure 2**, **Table S1**). Although high maternal proportions are also observed in the PE-like cross A×P (median of 0.769, the third-highest median; **Figure 2**), lower maternal proportions characterize the second PE-like cross C×P, falling within the range of those obtained for the weak-HSF crosses A×C and C×A that typically yield mixtures of viable and inviable seeds (Tukey-test, *P* <0.05; **Figure 2**; **Table S1**).

**Figure 2.**
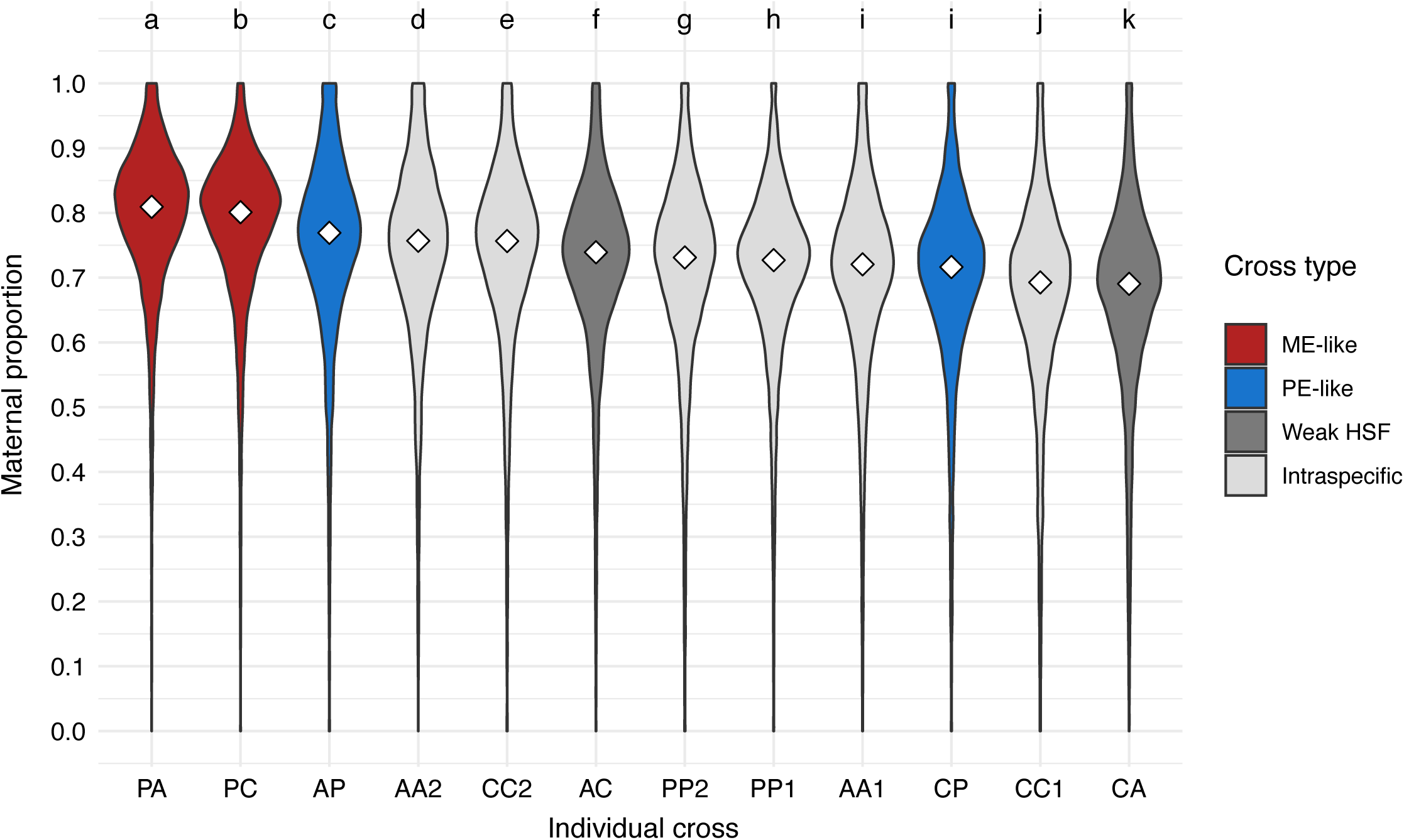
Violin plots representing the distribution of endosperm maternal proportions in all crosses within and between lineages A, C and P (ordered from left to right by decreasing median maternal proportion). Medians are indicated by open diamonds. ME-like, maternal-excess-like crosses; PE-like, paternal-excess-like crosses; Weak HSF, partial hybrid seed failure; Intraspecific, intraspecific (fully viable) crosses. Results from multiple comparisons of means are represented by letters at the top; distributions with significantly different means are indicated with different letters (Tukey’s test, *P* < 0.05). The number of informative genes for each cross is detailed in **Table S1**.

### Shifts in parental expression proportions in hybrid endosperms

To more precisely quantify (possibly systematic) changes in parental proportions upon hybridization, we estimated so-called ‘shifts’ in parental expression proportions at genome-wide scales. This necessitates two complementary comparisons, one with the maternal intraspecific reference data and another one with the paternal intraspecific reference data (**Dataset S2**). By estimating differences between expression proportions in hybrid endosperm and those in intraspecific endosperms sharing the same mother (‘shifts in maternal proportion’) and, separately, sharing the same father (‘shifts in paternal proportion’), we thus obtained two separate per-gene parental shift estimates that are mathematically independent. It is expected that maternal and paternal shifts are negatively correlated metrics. Regarding maternal shifts, the ME-like crosses P×A and P×C display marked genome-wide increases in maternal proportions (median increase of 0.08 and 0.07, respectively; **Figure 3**; **Table S1**). To a somewhat lesser extent, maternal proportions also display genome-wide increases in PE-like crosses A×P and C×P as well as in the A×C cross (median increases between 0.024 and 0.052; **Figure 3**), while the median maternal shift is close to zero in C×A (–0.003; **Table S1**). In terms of shifts in paternal expression, we found marked genome-wide decreases in paternal expression proportions for P×A, P×C and A×P (–0.047 to –0.031), in accordance with the notable elevated maternal expression proportions in these hybrid endosperms. However, data for C×P, C×A and A×C exhibit modest-to-high increases in paternal proportions (0.017, 0.070 and 0.018, respectively; **Figure 3**, **Table S1**) that are not easily reconcilable with “naïve” expectations, thus suggesting other factors at play (see **Discussion**).

**Figure 3.**
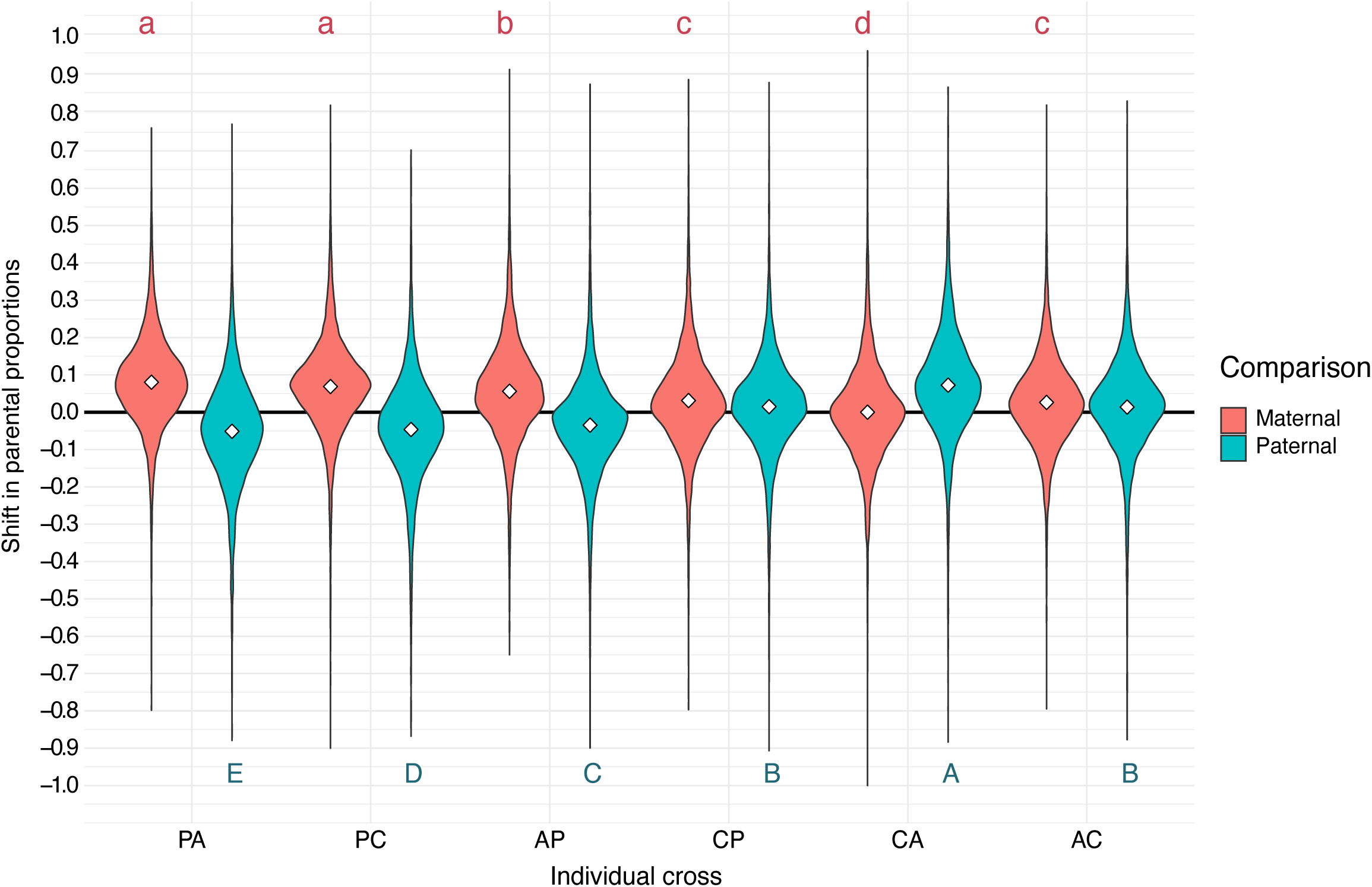
Distributions of shifts in parental expression proportions (hybrid–intraspecific reference) in the six hybrid crosses. Median values are indicated by open diamonds (cf. **Table S1**). Positive shift values imply higher parental proportion in hybrid endosperm compared to the intraspecific reference. For example, P×A is compared to PP1 as maternal reference (red) and to AA2 as paternal reference (turquoise). Crosses are ordered from left to right: ME-like (P×A and P×C), PE-like (A×P and C×P), and weak HSF (C×A and A×C). Tukey tests were performed independently for maternal and paternal comparisons, and significantly different distributions are indicated with different letters above (maternal) and below (paternal) the plotted data.

### Differential gene expression in the six unidirectional hybrid endosperms

We previously reported patterns of differential gene expression (DE) between hybrid and intraspecific endosperms where each hybrid cross was compared to pooled intraspecific data obtained from both parental species (Roth et al. 2019). To dissect possible parent-specific influences on DE in detail, we here perform a complementary analysis considering DE between each hybrid and (separately) both of the intraspecific endosperms sharing the same parents. For our six unidirectional hybrid crosses, the resulting 12 independent analyses reveal DE for 1.5 to 35.9% of the 22,006 tested genes (**Table S2**, **Dataset S3**). The overall pattern clearly indicates that gene expression in hybrid endosperms is more widely perturbed in comparison to their paternal intraspecific reference than in comparison to their maternal reference (Wilcoxon signed-rank test with H_1_ hypothesis “fewer DEs in maternal than in paternal comparisons”: *P* = 0.0156; **Dataset S3**). A second marked pattern reveals the ME-like endosperms to exhibit the highest proportions of DE genes, with larger fractions being underexpressed rather than overexpressed in hybrids, irrespective of whether comparisons consider the maternal or paternal reference data (54–62% of all DE genes in the four comparisons; **Table S2**).

Gene ontology (GO) analyses of differentially expressed genes in parental-excess-like hybrid endosperms revealed broad and coherent functional enrichments that are consistent with previously described transcriptional signatures in *Solanum* hybrids (Roth et al. 2019). Across both maternal and paternal reference comparisons, shared DE gene sets were predominantly enriched for processes related to chromatin organization and epigenetic regulation, transcriptional control (including genetic imprinting), DNA replication, and cell-cycle regulation, as well as core aspects of endosperm development and cellular proliferation (**Dataset S4**). Additional enrichments associated with hormone signaling, nutrient-reservoir activity, and proteolysis further indicate widespread perturbation of endosperm growth and metabolic functions. Taken together, these analyses support the conclusion that parental-excess-like hybrid endosperms experience extensive disruption of chromatin-based regulatory pathways and developmental programs, in line with earlier transcriptomic observations (Roth et al. 2019).

### Differential parent-specific expression (DPE) in relation to parental shifts

To further disentangle patterns of differential parental expression (DPE) in hybrid endosperms, we performed parent-specific differential gene expression analyses (**Figure 1**). Comparisons of parent-specific expression levels between hybrid and intraspecific endosperms yield two types of contrasts, depending on whether hybrid expression is compared to its maternal or to its paternal intraspecific reference data. A plausible null hypothesis posits that repression of the maternal allele ought to result in lower maternal expression proportion, and activation of the maternal allele ought to result in higher maternal proportion; the same rationale should apply for expression changes restricted to the paternal allele. Curiously, these predictions seem to have never been investigated in prior work. Our results are consistent with these expectations for DPE genes that are activated in hybrid endosperms: activation of the maternal allele coincides with distributions shifted toward more maternal expression proportions in hybrid endosperms (**Figure S2A**), and similarly, genes with upregulated paternal expression tend to show elevated paternal expression proportions. Consistent with these patterns, genes paternally repressed in hybrid endosperms show markedly negative paternal shifts (**Figure S2B**).

However, genes with repressed maternal expression do not exhibit the expected negative maternal shift in hybrid endosperms; this is most prominent in the ME-like endosperms where the majority of maternally repressed genes show positive maternal shifts (**Figure S2A**). This pattern suggests that in ME-like crosses, simultaneous paternal expression changes (whether or not significant) might have non-negligible impacts on parental ratios. In these crosses, DE gene underexpression might be caused by both paternal and maternal expression changes, and stronger reductions in paternal expression may lead to elevated maternal proportions.

### Asymmetric patterns of parent-specific expression perturbations

As total gene expression comprises both maternal and paternal transcripts, we next combined maternal- and paternal-specific DPE results for each hybrid class. For each gene, we classified expression changes depending on whether differential hybrid expression is limited to one parental allele (changes MAT-up/MAT-down and PAT-up/PAT-down), or affecting both parental contributions in the same direction (MATCH-up/MATCH-down) or in opposite directions (MAT-up/PAT-down and MAT-down/PAT-up; see Materials and Methods). In comparisons between parental expression proportions in hybrid endosperms and their intraspecific reference data, we found DPE in ∼8% to ∼29% of the common gene universe of 5,015 genes (**Table 1**, **Dataset S5**). Notably, ME-like hybrid endosperms exhibit larger numbers of DPE genes (29% and 24% of all informative genes in the P×A and P×C data, respectively) and higher proportions of DE among these DPE genes than other hybrid categories. Interestingly, paternal expression changes dominate in ME-like endosperms whereas maternal expression changes disproportionately dominate in PE-like endosperms (**Table 1**, **Dataset S5**). The ‘matching’ and ‘opposite’ parental expression changes are much less frequent in our data, at the exception of the ME-like endosperms P×A and P×C which encompass higher numbers of ‘matching’ expression changes (57 to 100 genes per category; **Table 1**, **Dataset S5**).

**Table 1.** Numbers of differentially parentally expressed (DPE) genes between intraspecific and hybrid endosperms, taking into account both parental (intraspecific) crosses sharing the same parents.

| DPE category | ME-like |  | PE-like |  | Weak HSF |  |
| --- | --- | --- | --- | --- | --- | --- |
|  | P×A–intra | P×C–intra | A×P–intra | C×P–intra | C×A–intra | A×C–intra |
| MAT ↓ | 190 | 212 | **344** | **156** | 65 | 180 |
| MAT ↑ | 176 | 157 | **346** | **180** | 139 | 236 |
| PAT ↓ | **451** | **332** | 26 | 11 | 92 | 54 |
| PAT ↑ | **446** | **341** | 73 | 37 | 117 | 87 |
| MATCH ↓↓ | 100 | 79 | 10 | 3 | 6 | 7 |
| MATCH ↑↑ | 69 | 57 | 15 | 7 | 8 | 8 |
| MAT ↓ PAT ↑ | 8 | 7 | 1 | 0 | 3 | 5 |
| MAT ↑ PAT ↓ | 9 | 6 | 3 | 0 | 2 | 2 |
| Sum of DPE genes | 1,449 | 1,191 | 818 | 394 | 432 | 579 |
| % DPE genes that are DE | 51.1 | 51.7 | 35.1 | 26.7 | 21.5 | 10.9 |

E.g. in P×A–intra, P×A is compared to PP1 for maternal expression and to AA2 for paternal expression. For all comparisons, the number of assessed genes is 5,015. Expression change categories are: MAT ↓ and MAT ↑, ‘repression’ or ‘activation’ restricted to the maternal allele (no significant change in expression of paternal allele); PAT ↓ and PAT ↑, repression or activation restricted to the paternal allele (no significant change in expression of maternal allele); MATCH ↓↓ and MATCH ↑↑, joint repression or activation of both parental alleles; MAT ↓ PAT ↑ and MAT ↑ PAT ↓, maternal and paternal alleles show opposite, significant expression changes. Numbers in bold highlight the most frequent categories of DPE changes involving those hybrid crosses resulting in near-complete hybrid seed failure (HSF). DE, differentially expressed.

### Contributions of DPE genes to shifts in hybrid parental proportions

To further understand how DPE may affect parent-specific expression as measured by maternal proportion, we binned DPE genes according to the magnitude of shifts in maternal proportion. Collectively, genes with parent-specific expression changes between hybrid and intraspecific endosperms contribute to both strongly negative and strongly positive shifts in maternal proportion (**Figure 4)**. This analysis reveals a gradual transition to more frequent repression of maternal and/or activation of paternal alleles coinciding with more negative shifts in maternal proportion, and vice versa, more frequent activation of maternal and/or repression of paternal alleles coinciding with more positive shifts in maternal proportion (**Figure 4**). Strikingly, and fully consistent with the strongly asymmetric DPE patterns summarized in **Table 1**, the contributions of maternal-specific changes to maternal shifts are particularly pronounced in the PE-like A×P and C×P endosperms (as well as in the more ‘balanced’ A×C endosperm), and the contributions of paternal-specific changes to maternal shifts are more pronounced in the ME-like P×A and P×C endosperms (as well as in the more ‘balanced’ C×A endosperm; **Figure 4**). Performing the equivalent analyses using estimates of paternal expression shifts yields corroborating evidence for these striking expression patterns (**Figure S3**).

**Figure 4.**
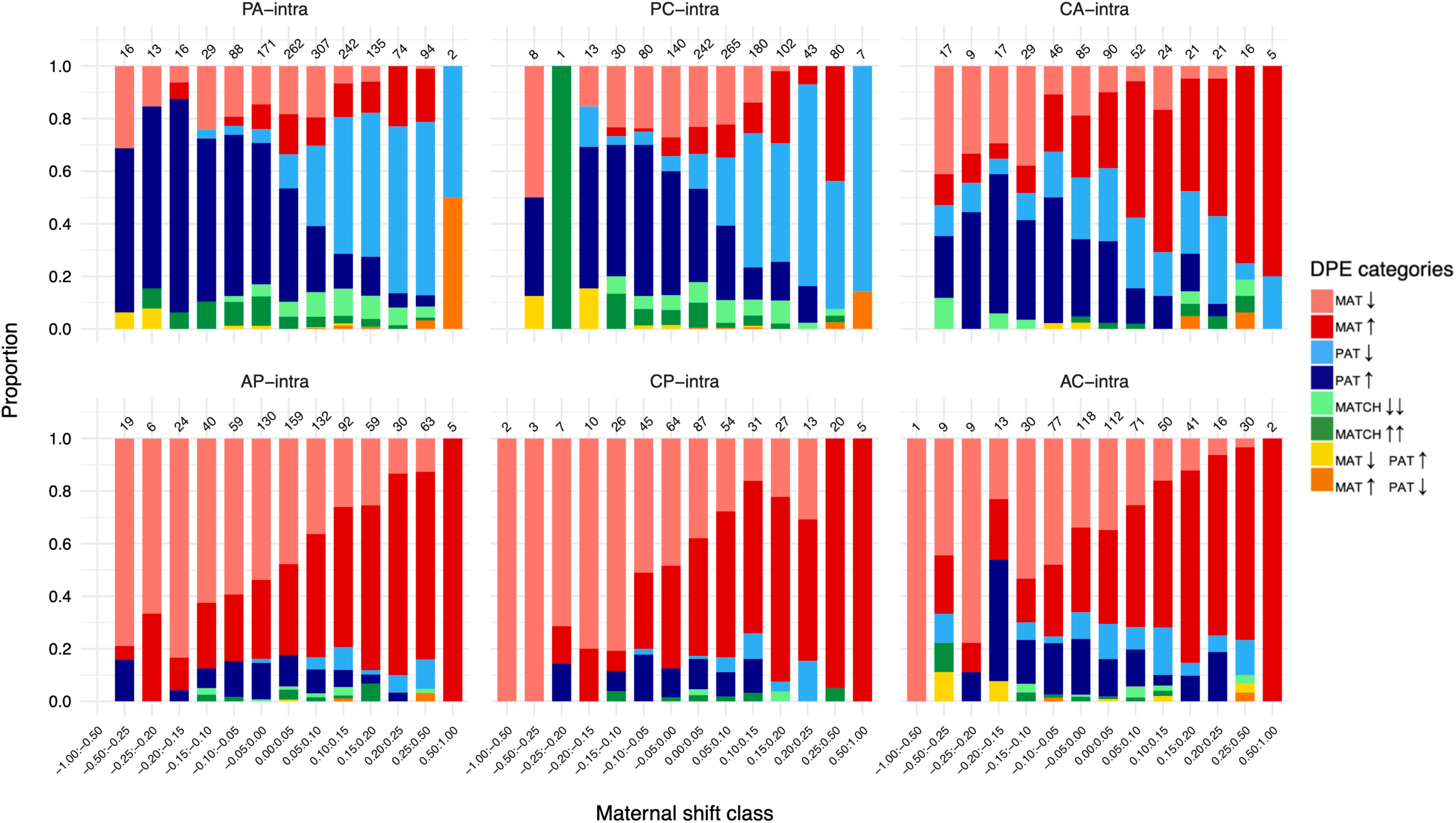
Maternal and paternal expression changes per maternal shift class in comparisons between hybrid endosperms and their corresponding intraspecific reference. Numbers at the top of the bars show the total number of DPE genes in each shift class (bin). A common set of 5,015 genes was assessed in each of the six comparisons and only genes with DPE are shown (cf. **Table 1**). Note that maternal shift classes between –0.25 and 0.25 have interval size 0.05 but that outside this range, intervals are larger, as indicated along the bottom x-axis.

### Cross-consistent parent-of-origin expression defines *Per*-associated gene classes

Next, we compiled the intersection of genes showing DPE in hybrid crosses that share the same parental-excess class. Here, cross-consistent refers to genes that show the same parent-of-origin misregulation direction across two related crosses (e.g. both ME-like crosses). Our ‘neutral’ expectations are derived from the numbers of DPE genes in **Table 1**, assuming that parent-specific misregulation categories occur independently between crosses, while conditioning on the total number of shared DPE genes. **Table 2** illustrates this concept with the ME-like crosses P×A and P×C, featuring 615 shared DPE genes. This comparison reveals two notable patterns. First, for each pairwise combination of DPE-direction categories between the two crosses (e.g. PAT-up or -down in both P×A and P×C), genes overwhelmingly fall into the same DPE category in both crosses. These concordant category combinations are highlighted in **Table 2** with gene numbers listed in bold italic font (Fisher’s exact test results in **Dataset S6**). Second, category combinations highlighted in orange or red are over-represented relative to independence expectations, whereas those in blue are under-represented (white indicates no clear deviation). Expected counts for the P×A–P×C comparison are shown in **Table S3**. Equivalent analyses for the PE-like crosses A×P and C×P are summarized in **Tables S4** and **S5**. Corresponding empirical and expected DPE-direction patterns for reciprocal cross pairs (C×P–P×C, A×P–P×A, and A×C–C×A) are provided in **Tables S6**-**S11**.

**Table 2.** Empirical contingency table of shared DPE genes between the ME-like crosses P×A and P×C.

|  |  | P×C |  |  |  |  |  |  |  |
| --- | --- | --- | --- | --- | --- | --- | --- | --- | --- |
| P×A | DPE category | MAT ↓ | MAT ↑ | PAT ↓ | PAT ↑ | MATCH ↓↓ | MATCH ↑↑ | MAT ↓ PAT ↑ | MAT ↑ PAT ↓ |
|  | MAT ↓ | **73** | 0 | **12** | 1 | **21** | 0 | 2 |  |
|  | MAT ↑ | 1 | **60** | 6 | **16** | 0 | **15** |  | 1 |
|  | PAT ↓ | 17 | 11 | **99** | 7 | **12** | 2 | 1 | 3 |
|  | PAT ↑ | 7 | 10 | 17 | **101** | 0 | **5** |  |  |
|  | MATCH ↓↓ | **31** | 0 | **11** | 0 | **24** | 0 |  |  |
|  | MATCH ↑↑ | 0 | **10** | 0 | **13** | 0 | **21** |  |  |
|  | MAT ↓ PAT ↑ | 1 |  |  |  |  |  |  |  |
|  | MAT ↑ PAT ↓ |  | 3 |  |  |  |  |  | 1 |

**Table 3.**
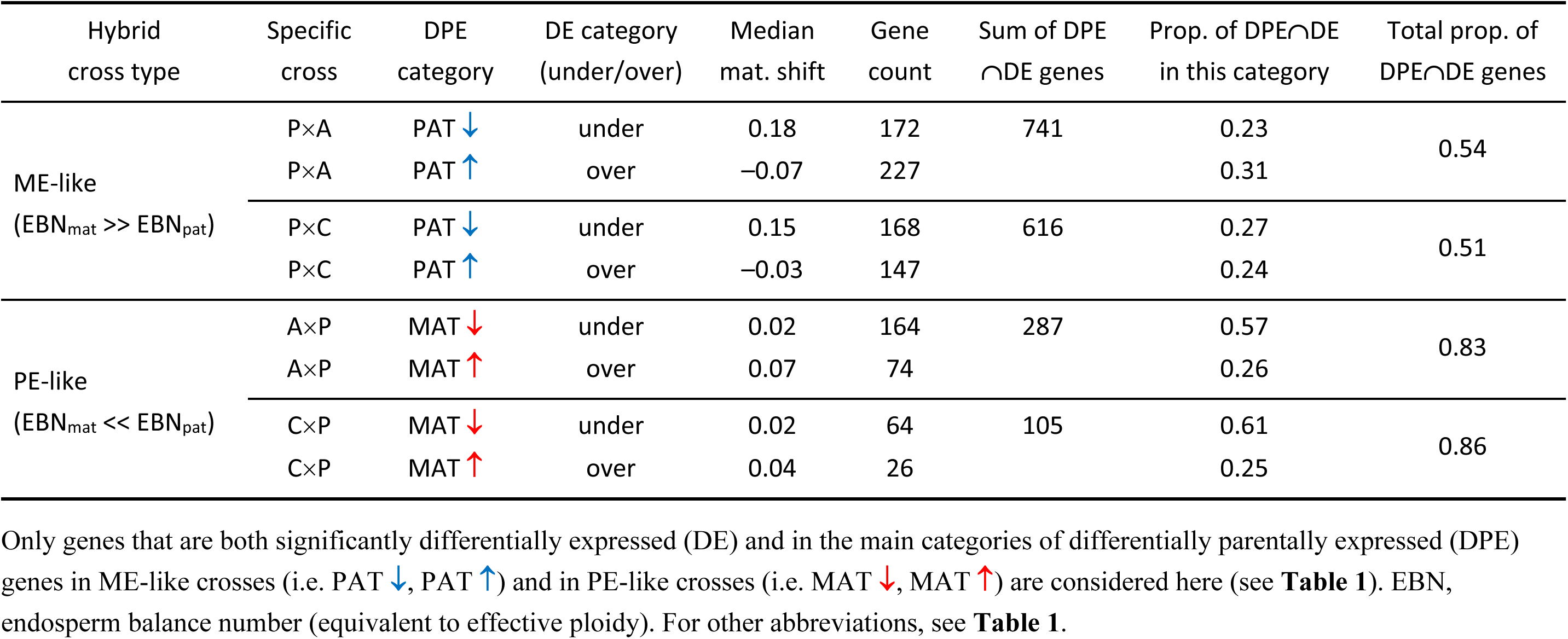
Contingency table corresponding to patterns of parent-specific expression relative to total expression and shifts in maternal (mat.) expression proportion.

| Hybrid cross type | Specific cross | DPE category | DE category (under/over) | Median mat. shift | Gene count | Sum of DPE ∩ DE genes | Prop. of DPE ∩ DE in this category | Total prop. of DPE ∩ DE genes |
| --- | --- | --- | --- | --- | --- | --- | --- | --- |
| ME-like<br>($EBN_{mat} \gg EBN_{pat}$) | P×A | PAT ↓ | under | 0.18 | 172 | 741 | 0.23 | 0.54 |
| | P×A | PAT ↑ | over | -0.07 | 227 | | 0.31 | |
| | P×C | PAT ↓ | under | 0.15 | 168 | 616 | 0.27 | 0.51 |
| | P×C | PAT ↑ | over | -0.03 | 147 | | 0.24 | |
| PE-like<br>($EBN_{mat} \ll EBN_{pat}$) | A×P | MAT ↓ | under | 0.02 | 164 | 287 | 0.57 | 0.83 |
| | A×P | MAT ↑ | over | 0.07 | 74 | | 0.26 | |
| | C×P | MAT ↓ | under | 0.02 | 64 | 105 | 0.61 | 0.86 |
| | C×P | MAT ↑ | over | 0.04 | 26 | | 0.25 | |

Summarizing these conspicuous patterns, our data reveal that although the ‘weaker’ parent exhibits higher numbers of DPEs in both parental roles (bold gene numbers in **Table 1**), higher proportions of DPE genes perturbed in the ‘stronger’ parent *Per* are shared in both ME-like and PE-like cross comparisons (red bars for P×A, P×C, and blue bars for A×P, C×P; **Figure 5**). Compared to the mean proportion of ‘DPE genes in the two contingency tables’ (grey bars), the opposite is true for the majority classes of DPE genes (perturbed in the weaker parent; blue bars for P×A, P×C, and red bars for A×P, C×P; **Figure 5**). Overall, *Per*-associated DPE genes are shared more consistently across ME-like and PE-like cross comparisons than DPE genes associated with the weaker parent. Notably, DPE genes in the MATCH-up/-down categories are also overrepresented in the two contingency tables, particularly in P×A and P×C (purple bars; **Figure 5**).

**Figure 5.**
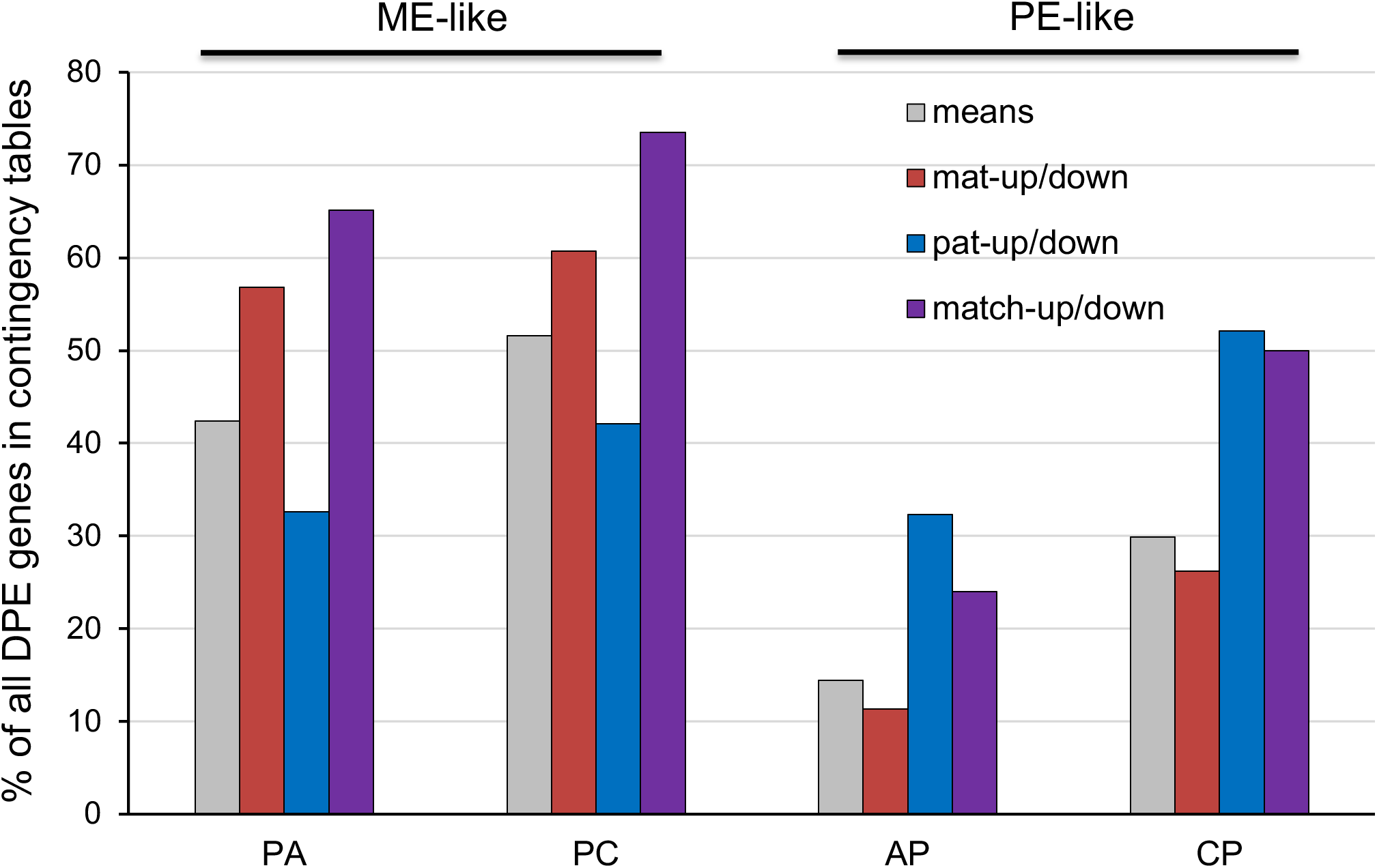
Proportions of all detected DPE genes (cf. **Table 1**) that are shared between the ME-like P×A and P×C hybrid categories (i.e. those that occur in their contingency table; **Table 2)**, as well as those shared between the PE-like A×P and C×P hybrid categories (**Table S4**). Breaking this down to different DPE categories reveals that higher proportions of the ‘minority DPE categories’ are shared in both ME-like (red bars) and PE-like hybrids (blue bars), compared to the ‘majority DPE categories’ (blue bars in ME-like and red bars in PE-like hybrids).

Analyses of the contingency tables for reciprocal hybrid crosses reveal that parent-specific expression perturbations seem to be mainly driven by effective-ploidy differences between the lineages and are not ‘parent-specific’ in the canonical sense of genomic imprinting. For example, 44 DPE genes in **Table S8** are MAT-down in A×P and PAT-down in the reciprocal cross P×A. In the same vein, a different suite of 73 DPE genes is MAT-up in A×P but PAT-up in the reciprocal cross P×A. In all such cases, the vast majority of DPE genes shows the same direction of parental misexpression in the independent hybrid crosses, either repression or activation. Together, these cross-context consistencies motivate a class-based summary of *Per*-associated DPE.

### Defining four *Per*-centered functional classes of DPE genes

Given these strong patterns, we here deduce four ‘functional classes’ of DPE genes, compiled across the four unidirectional hybrid crosses that comprise ME-like and PE-like data. For example, the 99 genes in **Table 2** that are PAT-down in both P×A and P×C are in the class “*repressed-by-Per*”, as are the 43 genes in **Table S4** that are MAT-down in both A×P and C×P. The 101 genes in **Table 2** that are PAT-up in both P×A and P×C are in the class “*activated-by-Per*”, as are the 73 genes in **Table S8** that are MAT-up in A×P and PAT-up in the reciprocal cross P×A. Continuing this rationale, the 73 genes in **Table 2** that are MAT-down in both P×A and P×C are in the class “*repression-of*-*Per*”, as are the 5 genes in **Table S4** that are PAT-down in both A×P and C×P. Finally, the 60 genes in **Table 2** that are MAT-up in both P×A and P×C illustrate the class “*activation-of-Per*”, as do the 15 genes in **Table S4** that are PAT-up in both A×P and C×P. We considered the proper combinations of ‘MATCH-up/-down’ and those with MAT-specific expression changes in the other cross (of which there are quite a lot in the P×A–P×C data (**Table 2**) but very few in other tables) to be part of *“activation-of-Per*” and “*repression-of-Per*”, respectively. The full data on DPE functional classes are compiled in **Dataset S7**. We next asked whether these *Per*-centered classes show coherent functional signatures.

### Functional signatures of DPE classes implicate chromatin regulation and hormone pathways

The “*repressed-by-Per*” category comprises 185 genes with conserved repression of the non-*Per* allele across reciprocal crosses **(Dataset S7)**. Several key chromatin regulators are found within this set, most notably Solyc01g094800, an SNF2-related chromodomain–helicase–DNA-binding protein, and Solyc07g063900, an RNA-binding protein, both of which show consistent repression of the non-*Per* allele across all four pairwise comparisons. Additionally, Solyc07g064090 (*FIE*), a core component of the Polycomb Repressive Complex 2 (PRC2), is found repressed in two comparisons. In STRING network analyses, these genes form a distinctive interaction module, centered around *FIE* and including Solyc04g081150 (Histone H3) and Solyc04g081350 (transcription factor E2F) (**Figure 6A)**. Together, these patterns identify a *Per*-associated repression module enriched for chromatin and transcriptional regulators and organized around a PRC2-centered interaction network. The possible implications of this coordinated repression for imprinting stability and HSF are discussed below.

**Figure 6.**
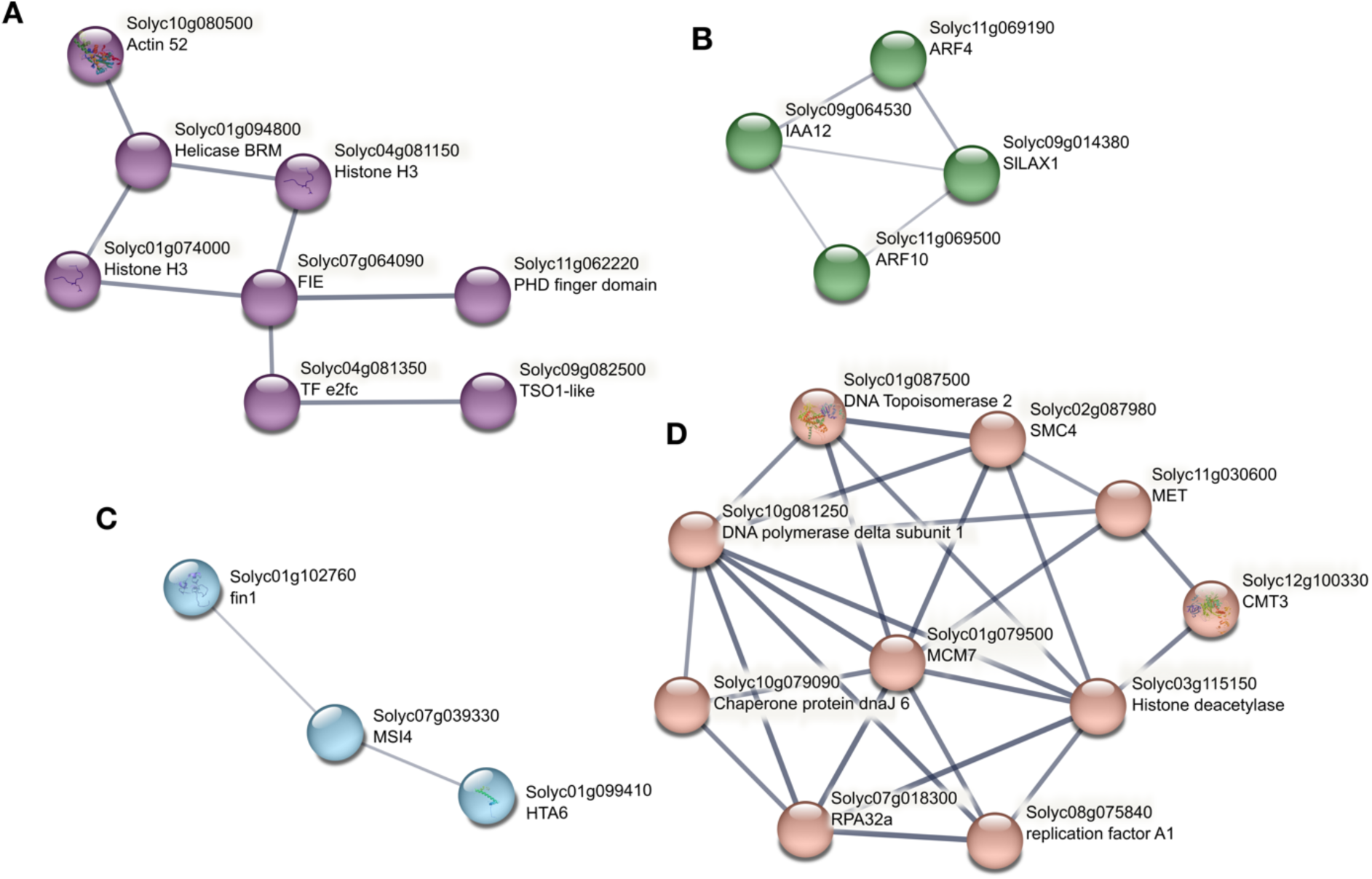
Proteins involved in epigenomic processes have distinct DPE patterns. **A.** Cluster of interacting proteins exemplifying the functional DPE class ‘*repressed-by-Per*’. The cluster centers on a core component of PRC2, FIE and includes two Histone H3 variants. **B.** Cluster of interacting proteins exemplifying the functional DPE class ‘*activated-by-Per*’, with auxin signaling functions. **C.** Cluster of interacting proteins exemplifying the functional DPE class ‘*activation-of-Per’* which are involved in chromatin organization, including a Histone H2 variant and a putative member of PRC2 in tomato, MSI4. **D.** Cluster of interacting proteins illustrating the functional DPE class ‘*repression-of-Per*’, including several proteins involved in the maintenance of methylation, replication, and chromatin organization.

The “*activated-by-Per*” category encompasses a set of 205 genes in which the non-*Per* allele is consistently activated across reciprocal hybrid crosses, independent of *Per*’s parental role (**Dataset S7)**. This recurrent pattern of DPE indicates that *Per* exerts a directional epigenetic effect, enhancing the expression of the alternate allele, regardless whether it is maternally or paternally inherited. Among the most striking components of this gene category is a cluster of auxin-related genes, whose expression is consistently activated by *Per* in hybrid endosperm **(Figure 6B)**. These include Solyc09g014380 (*SlLAX1*), an auxin influx carrier; Solyc11g069500 (*ARF10*) and Solyc11g069190 (*ARF4*), two auxin response factors; Solyc09g064530 (*IAA12*), an auxin/IAA transcriptional repressor; Solyc07g064280 (tryptophan synthase beta chain 1), involved in tryptophan biosynthesis and thus auxin production; and Solyc03g121270 (an IAA amino-acid hydrolase), which modulates auxin homeostasis. The coordinated activation of these genes suggests disruption of auxin-signaling dynamics, a pathway known to be critical for endosperm cellularization in *Arabidopsis* (Figueiredo et al. 2016). The enrichment of auxin-related regulators among *Per*-associated gene classes indicates that disruption of hormone-dependent developmental transitions is a recurrent and likely dosage-sensitive component of HSF.

The category “*activation-of-Per*” comprises 130 genes. Although functional enrichment is somewhat limited, one prominent interaction network involves Solyc11g073250, encoding a histone H2A protein, Solyc09g008850, an alfin-like transcription factor involved in chromatin remodeling, and Solyc02g083270, a putative chromatin-associated protein **(Figure 6C)**. In contrast, the “*repression-of-Per*” category reveals a more cohesive functional pattern **(Dataset S7)**. This group includes several central components of epigenetic regulation, most notably Solyc11g030600 (*MET1*) and Solyc12g100330 (*CMT3*), two methyl-transferases that in addition to Solyc11g007580 (*DEMETER*) drive the enrichment of GO:0044728 (DNA methylation or demethylation; *P* = 0.009). These genes, together with a cohort of replication-associated proteins—Solyc01g079500, Solyc03g115050, Solyc08g075840, and Solyc10g081250—also contribute to the enrichment of ‘DNA replication’ (KW-0235 ; *P* = 0.0012). Additional members such as Solyc03g115150 (a histone deacetylase), Solyc02g087980 (structural maintenance of chromosomes protein 4), and Solyc01g079500 (a DNA helicase of the MCM family), contribute to the enrichment of GO:0006325 (chromatin organization; *P* = 0.0365) and form a set of interacting proteins **(Figure 6D)**.

These findings suggest that the *Per* contribution of expression has a reduced core regulatory module of chromatin remodeling, replication, and methylation. Such repression likely compromises the establishment and maintenance of silencing marks, particularly those required for imprinting. This could explain the broader “*by-Per*” effects on gene expression, such as the activation of otherwise silenced alleles. Furthermore, the presence of the three DNA-directed RNA polymerases in this repressed set—Solyc06g053330, Solyc12g015780, and Solyc10g078860—raises the possibility that *Per* interferes with non-canonical RNA-directed DNA methylation (RdDM) pathways in tomato **(Dataset S7)**. Disruption of these pathways may lead to the observed reduction in small-RNA abundance (Florez-Rueda et al. 2021b) and the widespread failure of epigenetic silencing in hybrid seeds.

### Hybrid expression changes in candidate-imprinted genes

We explored expression changes upon hybridization within the set of conserved candidate-imprinted genes that we previously reported for the three focal lineages of wild tomatoes (Roth et al. 2018b). Regarding total expression changes, there is a striking pattern with DE maternally expressed genes (MEGs) being predominantly overexpressed and DE paternally expressed genes (PEGs) exclusively underexpressed across most types of hybrid endosperms (**Table S12**). In particular, a large number of candidate-imprinted genes exhibit DE in ME-like crosses P×A and P×C, with 54 to 59% of MEGs overexpressed compared to the maternal reference, and 29 to 53% of PEGs underexpressed compared to the paternal reference. For the PE-like crosses, 35% of the assessible PEGs are underexpressed in the A×P endosperm, yet none of the candidate-imprinted genes is DE in the C×P data. For the partly viable A×C and C×A crosses, we found only one PEG to be underexpressed in the C×A data (**Table S12**).

Regarding DPE analyses, 46 of the total 59 candidate-imprinted genes found to be conserved in imprinting status across all three wild tomato lineages (Roth et al. 2018b) could be retained for our present analyses. Overall, we uncovered DPE in up to 52% of conserved MEGs and in up to 62% of conserved PEGs. The largest proportions of DPE imprinted genes characterize the P×A and P×C data, mainly with father-specific changes. We found the paternal allele to be activated in 13 and 10 of the total 33 MEGs for P×A and P×C, respectively, and the paternal allele to be repressed in seven and four of the total 13 PEGs, respectively. Only a few candidate-imprinted genes show DPE among the remaining hybrid crosses, with one to four MEGs classified as MAT-up in each of these crosses (**Table S12**).

Regarding changes in parental proportions in hybrid endosperms, candidate PEGs exhibit large positive maternal shifts in the ME-like crosses, with increases in maternal proportion of 0.29 (P×A) and 0.33 (P×C), associated with (independently estimated) decreases in paternal proportion of 0.32 and 0.35 (**Table S12**). It thus appears that imprinted genes tend to be more perturbed in ME-like crosses compared to other cross types, with marked asymmetric expression responses in the two groups of imprinted genes: MEGs mostly overexpressed, PAT-up, with no perceptible shifts in parental proportions (lower than the genome-wide median values), and PEGs often underexpressed, PAT-down, underpinning large positive maternal shifts. These opposite patterns are consistent with the scenario depicted in **Figure 7**.

**Figure 7.**
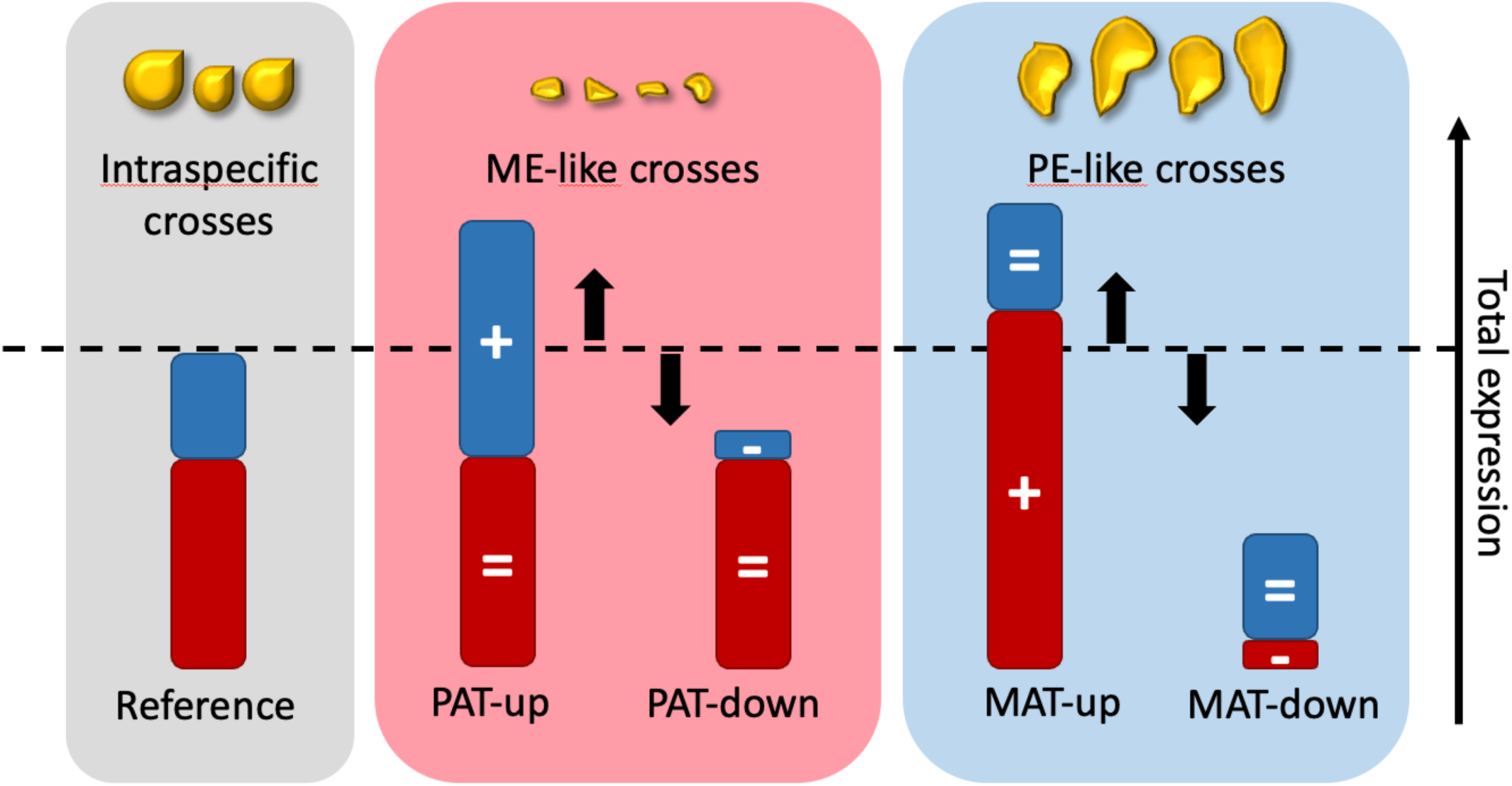
Qualitative scheme of the most frequently observed patterns of differential parental expression (DPE) in maternal-excess-like (ME-like) and paternal-excess-like (PE-like) crosses, compared to intraspecific, viable endosperms. PAT-up, PAT-down, MAT-up and MAT-down refer to the DPE categories summarized in **Table 1**. Typical mature seed phenotypes are shown at the top (Roth et al. 2018a; Städler et al. 2021). Red and blue bars represent maternal and paternal expression levels, respectively, whose sum yields total expression. Arrows and the +/-/= signs indicate the direction of change for total and parent-specific expression changes, respectively.

## DISCUSSION

### Genome-wide allelic polarization reflects lineage-specific *trans* regulation

Hybrid endosperm development is highly sensitive to divergence in parental regulatory states, yet it remains unclear how lineage differences translate into genome-wide shifts in allelic expression and their potential downstream consequences. By quantifying parent-of-origin expression across reciprocal wild-tomato crosses, we uncovered pervasive, cross-consistent polarization of parental contributions that tracks effective-ploidy differences between lineages. Our analyses reveal pronounced asymmetries in parent-specific expression patterns across hybrid endosperms, with the strongest and most consistent effects observed in crosses involving *Solanum peruvianum* (lineage *Per*). At a technical level, the higher number of genes amenable to parental expression analysis in *Per* reflects its higher level of nucleotide polymorphism (**Table S1**), but biologically, crosses involving this lineage repeatedly show the same directional biases in parental expression proportions across different hybrid combinations and reciprocal crosses. In other words, when *Per* participates in a cross, the shift in maternal versus paternal expression proportions tends to occur in a consistent direction across contexts, rather than varying idiosyncratically by partner species. In particular, ME-like crosses (in which *Per* is the ovule donor) display marked genome-wide increases in maternal expression proportions, accompanied by corresponding decreases in paternal contributions (**Figure 3**). By contrast, reciprocal and non-*Per* hybrid crosses show weaker or more heterogeneous patterns, including unexpected increases in paternal expression proportions in some cases (**Table S1**). Such cases may partly reflect within-lineage heterogeneity in effective ploidy, such that the specific *Chi*/*Ama* accessions sampled here could represent distributional “edge” genotypes with shifted parental-balance tendencies in particular pairings. Our finding of elevated maternal proportions even in crosses classified as PE-like, together with these non-intuitive paternal shifts, argues against a simple dosage-only model and instead points to lineage-specific regulatory dominance.

This interpretation is also consistent with genetic work in *Arabidopsis* showing that maternal:paternal transcript ratios in endosperm are actively regulated and can deviate from the naïve 2:1 expectation as part of a dosage-sensitive buffering system. In particular, the Pol IV/RdDM machinery has been implicated in maintaining global allelic balance, and perturbing this pathway can produce genome-wide shifts in parental contributions and strongly modify paternal-excess seed outcomes (Erdmann et al. 2017). Notably, Gehring and colleagues showed that the paternal state of canonical RdDM pathway genes is sufficient to sensitize paternal-excess seeds to paternal genome dosage, consistent with a model in which altered paternal epigenetic inputs trigger compensatory changes in allelic output—including increased maternal proportions—even under paternal-excess conditions (Satyaki et al. 2019). Moreover, parental Pol IV can exert distinct and even antagonistic parent-of-origin effects on endosperm gene expression, underscoring that parental contributions can be shaped by parent-specific epigenetic states established prior to fertilization rather than by dosage alone (Erdmann et al. 2017; Satyaki and Gehring 2022).

Importantly, such RdDM-associated buffering models predict coordinated allelic shifts across large gene sets rather than effects restricted to canonically imprinted loci, which aligns with our genome-wide DPE framework and the over-representation of MATCH-type coordinated allelic responses across reciprocal crosses. In our *Solanum* hybrids, several Pol IV/V-associated loci and sRNA-biogenesis components show reduced expression, and hybrid seeds exhibit globally reduced small-RNA abundance (Florez-Rueda et al. 2021b). This suggests that attenuation or rewiring of sRNA-directed methylation pathways may contribute directly to the observed, broad patterns of allelic rebalancing. Collectively, these results provide an external mechanistic framework in which lineage-associated divergence—here, the *Per* background—could engage conserved dosage-buffering architectures to generate repeatable, cross-context shifts in allelic balance. In this light, the disproportionate influence of *Per* on allelic expression balance in hybrid endosperms is consistent with its higher effective ploidy and previously reported dominance in both gene and small-RNA expression (Florez-Rueda et al. 2021b).

Importantly, the structure of parent-specific misexpression uncovered here does not conform to expectations from classical studies of genomic imprinting that predict stable parent-of-origin effects tied to maternal or paternal inheritance. Instead, reciprocal cross comparisons demonstrate that many alleles exhibit the same direction of misexpression regardless of maternal or paternal inheritance (see **Results**; **Figure 5**; **Tables 2, S4, S6-S11**). The over-representation of coordinated parental expression changes, including MATCH-up and MATCH-down categories (**Figure 5**), is consistent with *trans*-acting regulatory effects that would be expected to influence both alleles in the same direction (Wittkopp et al. 2008; Signor and Nuzhdin 2018). These patterns motivated our classification of DPE genes into four functional classes defined by their relationship to *Per* (**Dataset S7).** We next consider mechanistic entry points that might generate these *Per*-associated *trans*-allelic patterns.

### Mechanistic basis of *Per* dominance: *cis*×*trans* interactions and chromatin/small-RNA regulation

Among the *Per*-associated repression classes, ‘*repression-of-Per*’ highlights genes for which the *Per* allele is consistently repressed across cross contexts. This class is enriched for components of the epigenetic silencing machinery, including central DNA methyltransferases such as *MET1* and *CMT3*, histone deacetylases, RNA-directed DNA polymerases, and chromatin remodelers including members of the MCM complex, SNF2 family proteins, and CHD1-like helicases (**Figure 6D; Dataset S7**). The recurring involvement of these factors suggests that *Per*-associated allelic imbalance disproportionately affects pathways required for maintaining chromatin state and epigenetic homeostasis in hybrid endosperm. Within these classes, several signals converge on chromatin-based regulation, including Polycomb-linked repression, DNA methylation/RdDM components, and histone-variant dynamics.

In contrast, the ‘*repressed-by-Per*’ class captures genes whose non-*Per* alleles are consistently repressed in *Per*-containing hybrid contexts. Notably, this class includes FIE, a core component of the FIS-PRC2 complex and a central node in a chromatin-regulator interaction network (**Figure 6A**). Given the established role of PRC2 in imprinting control and endosperm development, including maintenance of repressive chromatin states at dosage-sensitive loci and PEG regulatory targets (Köhler et al. 2003; Gehring and Satyaki 2017; Batista and Köhler 2020), *Per*-associated repression within a PRC2-centered module raises the possibility that Polycomb-mediated silencing capacity is partially compromised in these hybrids. While our data do not directly measure PRC2 activity, reduced expression of PRC2-linked regulators could plausibly weaken repression fidelity at subsets of dosage-sensitive loci, contributing to the imprinting perturbations and allelic imbalance observed (Florez-Rueda et al. 2016; Roth et al. 2019). Importantly, in our dataset, genome-wide shifts in maternal proportion can be accompanied by reduced paternal contributions (**Figure 3**), implying that paternal downregulation and impaired maternal-allele silencing should be viewed as non-exclusive mechanisms. Consistent with this hypothesis framework, disruption of FIS-PRC2 components in *Arabidopsis* is associated with endosperm overproliferation, delayed or failed cellularization, and seed abortion, and PRC2 function is tightly linked to parental genome dosage and hybridization barriers (Baroux et al. 2007; Kradolfer et al. 2013; Lafon-Placette and Köhler 2016).

Concurrently, *Per* activates auxin-related and cell-cycle regulatory pathways in the non-*Per* alleles, further contributing to developmental instability. The ‘*activated-by-Per*’ gene set includes several key auxin-signaling components—*ARF10*, *ARF4*, *IAA12*, and *SlLAX1*—(**Figure 6B**), suggesting a shift toward auxin overproduction or misregulation. Since auxin regulates endosperm cellularization, the timing of growth transitions, and maternal control of nutrient allocation, its misregulation in hybrids may delay cellularization and/or cell proliferation, promote excessive nuclear proliferation, or disrupt nutrient partitioning. These results are consistent with previous findings implicating auxin imbalance in interploidy and interspecies seed failure in plants (Figueiredo and Köhler 2018; Florez-Rueda et al. 2025).

Consistent with a broader chromatin-based disruption suggested by the *Per*-associated classes, we also detect *Per*-linked DPE among histone-variant genes that can shape nucleosome stability and nuclear organization. In plants, variant-specific nucleosome composition helps define distinct chromatin environments—ranging from transcriptionally permissive to stably repressed domains—and is particularly relevant during the extensive chromatin reprogramming that accompanies reproduction and early seed development (Jiang et al. 2017; Borg et al. 2020; Foroozani et al. 2022; Jamge et al. 2023; Simon and Probst 2024). In our dataset, we identified several histone-variant DPE genes in hybrid endosperms, pointing to potential disruptions in chromatin organization as a consequence of *Per* dominance. Notably, Solyc03g005220 (H2AXb) was found in the ‘*repression-of-Per*’ category, and Solyc01g099410 (HTA6) and Solyc11g073250 (H2A isoform x1) were found among genes in the ‘*activation-of-Per*’ category. These observations are mechanistically plausible, given that H2A variants can influence DNA accessibility, nucleosome stability, and genome maintenance pathways. For example, in *Arabidopsis*, H2A.X contributes to maintaining DNA methylation homeostasis in the endosperm chromatin environment (Frost et al. 2023).

More broadly, disruption of heterochromatin organization has been directly linked to hybrid endosperm failure in *Capsella*, where reduced chromatin condensation and associated chromosome instability accompany seed arrest (Dziasek et al. 2021). In this context, *Per*-associated misregulation of histone-variant genes could plausibly contribute to altered higher-order chromatin organization and impaired chromosome-scale regulation in hybrid endosperm, potentially affecting how parental chromosome sets are functionally integrated during development. This histone-variant misregulation coincides with repression of chromatin remodeling and epigenetic silencing factors described above, consistent with broader perturbation of chromatin-state control in hybrid endosperm. Of particular interest here are two H3 variant genes—Solyc04g081150 and Solyc01g074000. H3 variant composition can modulate Polycomb-associated reprogramming during reproduction (Borg et al. 2020, 2021). Their repression in *Per*-associated contexts is therefore consistent with altered histone-based epigenetic memory or replacement dynamics, which could impair chromatin reprogramming of the paternal genome in hybrid endosperm. Given that the endosperm represents a central arena of parental dosage interaction and resource-allocation conflict, and that chromatin-state regulation contributes to this balance (Jiang et al. 2017; Simon and Probst 2024), altered histone-variant expression may contribute to endosperm malfunction.

Taken together, the four *Per*-centered DPE classes also point to the likely provenance of regulatory perturbation in hybrid endosperm. The “*by-Per*” classes (i.e. *activated-by-Per*; *repressed-by-Per*) capture systematic expression shifts in the accompanying non-*Per* allele in *Per*-containing hybrids and therefore most directly reflect a *Per*-associated postzygotic *trans*-acting regulatory context. In contrast, the “*of-Per*” classes (i.e. *activation-of-Per*; *repression-of-Per*) comprise consistent expression changes in the *Per* allele itself across cross contexts, indicating that the *trans*-dominant lineage is also responsive to the hybrid environment—consistent with *cis*-by-*trans* mismatch, in which *Per* allele-specific *cis*-regulatory architecture is interpreted in a divergent hybrid *trans*-regulatory context, yielding systematic *Per*-allele up- or down-shifts, and with compensatory or dosage-control responses that buffer developmental programs under genomic imbalance (Wittkopp et al. 2008; Erdmann et al. 2017; Signor and Nuzhdin 2018; Satyaki et al. 2019; Marand et al. 2023), rather than a purely unidirectional model in which *Per* simply imposes expression states on the partner genome. Together, these patterns are compatible with a model in which parental carry-over effects (maternal central-cell chromatin state and PRC2-mediated set-up; parent-of-origin small-RNA/RdDM inputs) interact with post-fertilization *trans* regulation to shape genome-wide allelic output in the endosperm (Erdmann et al. 2017; Köhler et al. 2021; Satyaki and Gehring 2022). Under this interpretation, *Per* dominance and *Per*-allele sensitivity are not contradictory but represent complementary signatures of a (higher effective ploidy) lineage that both imposes and reacts to perturbed regulatory landscapes in hybrid seeds.

### A *trans*-allelic dominance framework for hybrid seed failure

Together, the coordinated repression of both histone-variant genes and components of the silencing machinery points to a broader disturbance of chromatin-based regulatory control under *Per* dominance. If histone replacement dynamics and Polycomb-mediated repression are jointly weakened, as our data suggest for *Solanum*, this could prevent proper imprinting establishment, destabilize gene expression domains, and hinder the developmental transitions required for endosperm proliferation and seed viability. While these mechanisms remain to be tested directly, they offer a chromatin-level hypothesis linking *Per*-associated *trans*-dominant effects with epigenetic imbalance in hybrid endosperm.

Jointly, our findings support a model of *trans*-allelic epigenetic dominance, in which regulatory factors associated with *Per* reshape allelic expression landscapes in hybrid endosperm independently of parental origin, providing a mechanistic link between divergence in effective ploidy, widespread transcriptional imbalance, and HSF. More broadly, chromatin features and transcription-factor networks are likely to intersect with epigenetic methylation pathways and imprinted gene regulation, such that parent-of-origin biases in gene expression emerge from the combined effects of DNA methylation, histone modifications, nucleosome accessibility, and regulatory-factor binding that collectively define permissive and repressive chromatin states at key developmental loci (Gehring and Satyaki 2017; Batista and Köhler 2020; Marand et al. 2023; Simon and Probst 2024). In this context, reciprocal hybrid analyses that integrate chromatin accessibility and chromatin state profiling—together with methylome and small-RNA measurements—will be essential to disentangle parent-specific imprinting effects from broader *trans*-acting shifts and to resolve how lineage-specific *trans* regulation and parent-of-origin epigenetic asymmetries jointly shape endosperm developmental trajectories in hybrid seeds.

## MATERIALS AND METHODS

### Plant material and crossing design

Plant origins, maintenance, and the crossing design have been previously described in detail (Roth et al. 2018b, 2019), and population-genomic context for the Arcanum group is provided in Florez-Rueda et al. (2021a). Seeds were generously provided by the Tomato Genetics Resource Center at the University of California, Davis (https://tgrc.ucdavis.edu). In brief, the crossing design involved two plants of *Solanum arcanum* var marañón [LA1626 (Ancash, Peru) and LA2185 (Amazonas, Peru)], two plants of *S. chilense* [LA2748 (Tarapaca, Chile) and LA4329 (Antofagasta, Chile)], and two plants of *S. peruvianum* [LA2744 (Arica and Parinacota, Chile) and LA2964 (Tacna, Peru)]. We use lower-case letters to identify individual genotypes of these accessions, directly following the four-number LA accession numbers. These six genotypes were used for three reciprocal intraspecific crosses and three reciprocal hybrid crosses (the latter involving only LA2185a, LA4329b and LA2744b). We refer to reciprocal crosses with the two initial letters of parental lineages within brackets (reciprocal crosses are [AC], [AP], [CP], [AA], [CC] and [PP]), and to individual (i.e. unidirectional) crosses with the initial letters of parental lineages without brackets, indicating the cross direction ‘mother×father’: **AA1**, LA2185a×LA1626b; **AA2**, LA1626b×LA2185a; **CC1**, LA4329b×LA2748b; **CC2**, LA2748b×LA4329b; **PP1**, LA2744b×LA2964a; **PP2**, LA2964a×LA2744b; **A**×**C**, LA2185a×LA4329b; **C**×**A**, LA4329b×LA2185a; **A**×**P**, LA2185a×LA2744b; **P**×**A**, LA2744b×LA2185a; **C**×**P**, LA4329b×LA2744b; **P**×**C**, LA2744b×LA4329b. Based on our crossing design and adopted nomenclature, this implies that A×C, A×P and AA1 share the same mother, that C×A, C×P and CC1 share the same mother, and that P×A, P×C and PP1 share the same mother.

### Data generation

Endosperm sampling, library preparation and sequencing was performed as previously described (Roth et al. 2018b, 2019) and replicated three times for each cross (using three independent ramets grown from cuttings of each genotype), leading to a total of 36 laser-captured endosperm samples.

### Quantification of parent-specific expression proportions

Quality control, alignment to the SL2.50 tomato reference genome (The Tomato Genome Consortium 2012) and SNP-calling of RNA-seq data was performed as previously described (Roth et al. 2018b, 2019). Parental counts per gene were calculated using parental flower-bud transcriptome data and a custom python pipeline for genes polymorphic in the parental plants and with sufficient coverage (see Roth et al. 2018b). Sequence reads from the three replicates were combined for each cross before estimating parent-specific gene expression, with effects analogous to implementing mean estimates weighted by the replicates’ sequencing depth at polymorphic sites. The number of genes recovered for parent-specific gene expression estimates (i.e. those with parental sequence differences) per reciprocal cross is 13,198 for [PP], 9,623 for [CC], 7,730 for [AA], 11,195 for [AC], 11,414 for [AP] and 11,985 for [CP]. When counts were used for Differential Parental Expression (DPE) analysis, corrected parental counts for each gene were obtained from the parental allelic counts divided by the number of polymorphic sites. This allowed us to normalize counts with respect to different polymorphism levels across endosperm samples. When counts were used for visual plotting, a normalization step was added using the EdgeR function ‘calcNormFactors’ with the method ‘TMM’ (Robinson et al. 2010). For total expression levels, the transcript-per-million (TPM) values were obtained as previously described (Roth et al. 2018b).

### Shifts in parental expression proportions in hybrid endosperms

In each cross and for every gene with parental sequence differences (i.e. those with power to inform maternal vs. paternal expression proportions), maternal proportions were defined as the ratio between maternal and total (i.e. maternal + paternal) counts. For all hybrid endosperms, we calculated so-called ‘shifts’ in parental proportions: shifts in maternal proportions were calculated as the difference between the maternal proportions in the hybrid and its intraspecific reference cross sharing the same ovule parent, corresponding to differences referred to in the text as “P×A–PP1”, “A×P–AA1”, “P×C–PP1”, “C×P–CC1”, “C×A–CC1” and “A×C–AA1”. Correspondingly, shifts in paternal proportions were calculated as the difference between the paternal proportions (equal to 1 – maternal proportions) in the hybrid and its intraspecific reference cross sharing the same pollen parent, corresponding to differences referred to in the text as “P×A–AA2”, “A×P–PP2”, “P×C–CC2”, “C×P–PP2”, “C×A–AA2” and “A×C–CC2”.

### Analyses of differential expression (DE and DPE) in hybrid endosperms

For Differential gene Expression (DE) analyses, the gene universe comprises 22,006 genes; we only kept genes with at least one read count per million in at least two of the 36 libraries. Both DE and DPE analyses were performed with the R package EdgeR (Robinson et al. 2010; R Development Core Team 2014). For DPE analyses, one sample was removed (replicate 1 of AA2) and the gene universe is necessarily restricted to genes that are expressed and polymorphic (i.e. with parental sequence differences) in all remaining samples (35 libraries, 5,015 genes). Owing to these filtering steps, 33 of the 42 candidate Maternally Expressed Genes (MEGs) and 13 of the 17 candidate Paternally Expressed Genes (PEGs) identified as conserved imprinted genes across the three focal wild tomato lineages (Roth et al. 2018b) could be retained for DPE analyses; for DE analyses, all candidate imprinted genes except for one MEG were retained after filtering.

For DPE analyses, we performed separate analyses for maternal and paternal expression levels, contingent on the relevant intraspecific reference (see above). We fitted a negative binomial model to each gene using individual crosses as a factor, estimating trended dispersions (variance parameters). For each cross comparison, DE and DPE genes were identified with a generalized linear model and a quasi-likelihood *F*-test (Alessandrì et al. 2019). *P*-value correction was implemented after testing with the Benjamini–Hochberg method (Benjamini & Hochberg 1995) for a False Discovery Rate (FDR) of 5%. These comparisons were performed between hybrid and intraspecific endosperms sharing the same parents (P×A–intra, A×P–intra, P×C–intra, C×P–intra, C×A–intra and A×C–intra comparisons), employing two independent DPE tests to identify maternally and paternally ‘activated’ or ‘repressed genes’, taking into account the two parental intraspecific references. For instance, the cross P×A shares its mother with PP1 (plant LA2744b) and its father with AA2 (plant LA2185a); thus in the contrast “P×A–intra”, we compared the maternal counts of the P×A data to the maternal counts of PP1 and the paternal counts of the P×A data to the paternal counts of AA2. We report details on all tested comparisons for DE and DPE analyses in **Dataset S5.** We emphasize that no particular molecular mechanisms are invoked by our nomenclature; we simply use ‘activated’ and ‘repressed’ for DPE phenomena to distinguish them from the DE-associated terms ‘overexpressed’ and ‘underexpressed’.

### Categorization of genes with parent-specific activation or repression

To summarize and gauge parental expression changes via contrasting the results of DPE analyses performed on the same genes for maternal and paternal expression, we defined eight different expression categories. Expression changes significant for only one parental allele were classified as ‘MAT-up’ or ‘MAT-down’ when the maternal allele was found to be activated or repressed in hybrid endosperm, with no significant change in expression of the paternal allele, and as ‘PAT-up’ or ‘PAT-down’ when the paternal allele was found to be activated or repressed in hybrid endosperm, with no significant change in expression of the maternal allele. The categories ‘MATCH-up’ and ‘MATCH-down’ refer to joint activation or repression of both parental alleles in hybrid endosperm. Lastly, in the ‘MAT-up & PAT-down’ and ‘MAT-down & PAT-up’ categories, maternal and paternal alleles show opposite expression changes.

As patterns of conserved DPE across reciprocal crosses emerged, we constructed contingency tables for all relevant pairwise comparisons to identify genes consistently classified as “*repressed-by-Per*”, “*activated-by-Per*”, “*activation-of-Per*”, or “*repression-of-Per*”. For each comparison, we calculated the expected distribution of DPE combinations under the assumption of independent misregulation between crosses. The observed gene counts were then tested against these expectations using Fisher’s exact test, allowing us to identify categories that were significantly over-represented. Genes falling into these enriched categories are highlighted in bold italic font in the corresponding tables.

To detect enriched functions, analyses were performed in STRING v 11.5 (Szklarczyk et al. 2017). The gene universe was all the genes expressed. All reported enriched terms are significant with an FDR-corrected *P* <0.05. To delimit genes with discrete functions and plot protein networks, an MCL clustering algorithm with an inflation parameter of 3 was implemented in STRING. We used topGO to identify enriched Gene Ontology (GO) terms (Alexa and Rahnenführer 2016). The gene universe considered comprises the 5,015 genes used in the present DPE analyses.

## DATA AVAILABILITY

Raw RNA-seq data used in this study are deposited in the NCBI Sequence Read Archive under accessions **PRJNA427095**, **SRP132466**, and **SRX1850236** (previously published in Roth et al. 2018b, 2019).

## ACKNOWLEDGMENTS

We are grateful Tadeas Priklopil for advice on statistics, and Alex Widmer for his general support of this project. We thank the C.M. Rick Tomato Genetics Resource Center at U.C. Davis for supplying seed samples. This work was supported by the Swiss National Science Foundation [31003A_130702 to T.S.] and an ETH Research Grant [ETH-40 13-2 to T.S. and Alex Widmer].

---

**Table S1.** Overview of parental contributions to gene expression in both intraspecific and hybrid crosses.

| Unidirectional cross<br>(mother×father) | Cross type | Median maternal<br>proportion | No. of genes<br>assessable for<br>maternal prop. | Median shift in<br>maternal prop.<br>(*n* genes) | Median shift in<br>paternal prop.<br>(*n* genes) |
| --- | --- | --- | --- | --- | --- |
| AA1 | Intra-specific | 0.720 | 7,730 | – | – |
| AA2 | Intra-specific | 0.757 | 7,730 | – | – |
| CC1 | Intra-specific | 0.693 | 9,623 | – | – |
| CC2 | Intra-specific | 0.757 | 9,623 | – | – |
| PP1 | Intra-specific | 0.727 | 13,198 | – | – |
| PP2 | Intra-specific | 0.731 | 13,198 | – | – |
| P×A | ME-like | 0.809 | 11,414 | 0.078 (10,730) | -0.047 (7,243) |
| P×C | ME-like | 0.801 | 11,985 | 0.070 (11,245) | -0.041 (9,022) |
| A×P | PE-like | 0.769 | 11,414 | 0.052 (7,243) | -0.031 (10,730) |
| C×P | PE-like | 0.716 | 11,985 | 0.026 (9,022) | 0.017 (11,245) |
| C×A | Weak HSF | 0.691 | 11,195 | -0.003 (8,789) | 0.070 (7,156) |
| A×C | Weak HSF | 0.739 | 11,195 | 0.024 (7,156) | 0.018 (8,789) |

Median maternal expression proportions in each cross (column 3) and magnitude of shifts in parental proportions in hybrid endosperms (rightmost columns). –, not defined (no hybrid cross); HSF, hybrid seed failure; ME-like, maternal-excess-like; PE-like, paternal-excess-like (the latter two cross types result in near-complete HSF). The number of genes assessable for parental shifts (maternal or paternal, in parentheses) is lower than the number of genes assessed for maternal proportions as the former must be polymorphic both in the hybrid and the corresponding intraspecific cross. Differences in maternal proportions were calculated by comparing maternal proportions in A×P and A×C to AA1, in C×P and C×A to CC1, and in P×A and P×C to PP1, respectively. Differences in paternal proportions were calculated by comparing paternal proportions (1 – maternal proportion) in A×P and C×P to PP2, in C×A and P×A to AA2, and in A×C and P×C to CC2, respectively (see Materials and Methods).

**Table S2.** Statistics of differentially expressed (DE) genes in hybrid–intraspecific comparisons. Hybrid cross type.

| Hybrid cross type | Comparison<br>hybrid–intra | Underexpressed<br>in hybrid | Overexpressed<br>in hybrid | Sum of<br>DE genes |
| --- | --- | --- | --- | --- |
| ME-like<br>(EBN<sub>mat</sub> >> EBN<sub>pat</sub>) | P×A–PP1 | 2,900 | 1,766 | 4,666 |
|  | P×A–AA2 | 4,241 | 3,649 | 7,890 |
|  | P×C–PP1 | 2,019 | 1,401 | 3,420 |
|  | P×C–CC2 | 3,759 | 2,906 | 6,665 |
| PE-like<br>(EBN<sub>mat</sub> << EBN<sub>pat</sub>) | A×P–AA1 | 1,675 | 1,502 | 3,177 |
|  | A×P–PP2 | 1,884 | 1,858 | 3,742 |
|  | C×P–CC1 | 771 | 721 | 1,492 |
|  | C×P–PP2 | 1,016 | 1,120 | 2,136 |
| Weak HSF<br>(EBN<sub>mat</sub> +/- EBN<sub>pat</sub>) | C×A–CC1 | 134 | 201 | 335 |
|  | C×A–AA2 | 2,319 | 2,470 | 4,789 |
|  | A×C–AA1 | 267 | 396 | 663 |
|  | A×C–CC2 | 947 | 1,010 | 1,957 |

Expression data from each of six classes of hybrid endosperms were compared to their intraspecific maternal (red) and paternal reference (blue) expression levels; 22,006 genes were tested in each comparison. Maternal-excess (ME)-like and paternal-excess (PE)-like crosses result in very high proportions of inviable seeds, while ‘weak HSF’ refers to hybrid crosses yielding mixtures of viable and inviable seeds. EBN, endosperm balance number. Complete results of the DE analyses are in Dataset 5

**Table S3.** ***Expected numbers* of shared DPE genes between the ME-like crosses P×A and P×C, given the observed numbers in Table 1**.

|  |  | P×C |  |  |  |  |  |  |  |
| --- | --- | --- | --- | --- | --- | --- | --- | --- | --- |
|  | DPE category | MAT ↓ | MAT ↑ | PAT ↓ | PAT ↑ | MATCH ↓↓ | MATCH ↑↑ | MAT ↓ PAT ↑ | MAT ↑ PAT ↓ |
| **P×A** | MAT ↓ | 14.4 | 10.6 | 22.5 | 23.1 | 5.3 | 3.9 | 0.5 | 0.4 |
|  | MAT ↑ | 13.3 | 9.8 | 20.8 | 21.4 | 5.0 | 3.6 | 0.4 | 0.4 |
|  | PAT ↓ | 34.1 | 25.2 | 53.4 | 54.8 | 12.7 | 9.2 | 1.1 | 1.0 |
|  | PAT ↑ | 33.7 | 25.0 | 52.8 | 54.2 | 12.6 | 9.1 | 1.1 | 1.0 |
|  | MATCH ↓↓ | 7.6 | 5.6 | 11.8 | 12.2 | 2.8 | 2.0 | 0.2 | 0.2 |
|  | MATCH ↑↑ | 5.2 | 3.9 | 8.2 | 8.4 | 1.9 | 1.4 | 0.2 | 0.1 |
|  | MAT ↓ PAT ↑ | 0.6 | 0.4 | 0.9 | 1.0 | 0.2 | 0.2 | 0.0 | 0.0 |
|  | MAT ↑ PAT ↓ | 0.7 | 0.5 | 1.1 | 1.1 | 0.3 | 0.2 | 0.0 | 0.0 |

**Table S4.** Empirical contingency table of shared DPE genes between the PE-like crosses A×P and C×P.

|  |  | C×P |  |  |  |  |  |  |  |
| --- | --- | --- | --- | --- | --- | --- | --- | --- | --- |
|  | DPE category | MAT ↓ | MAT ↑ | PAT ↓ | PAT ↑ | MATCH ↓↓ | MATCH ↑↑ | MAT ↓ PAT ↑ | MAT ↑ PAT ↓ |
| A×P | MAT ↓ | **43** | 3 | 0 | 1 |  |  |  |  |
|  | MAT ↑ | 7 | **23** | 0 | 1 |  |  |  |  |
|  | PAT ↓ | 2 |  | 5 | 0 | 0 | 0 |  |  |
|  | PAT ↑ | 1 | 5 | 0 | **15** | 0 | 4 |  |  |
|  | MATCH ↓↓ | 1 |  | 0 | 0 | 1 |  |  |  |
|  | MATCH ↑↑ |  | 2 | 0 | 2 |  |  |  |  |
|  | MAT ↓ PAT ↑ |  | 1 |  |  |  |  |  |  |
|  | MAT ↑ PAT ↓ |  |  | 1 |  |  |  |  |  |

**Table S5.** ***Expected numbers* of shared DPE genes between the PE-like crosses A×P and C×P, given the observed numbers in Table 1**.

|  |  | C×P |  |  |  |  |  |  |  |
| --- | --- | --- | --- | --- | --- | --- | --- | --- | --- |
|  | DPE category | MAT ↓ | MAT ↑ | PAT ↓ | PAT ↑ | MATCH ↓↓ | MATCH ↑↑ | MAT ↓ PAT ↑ | MAT ↑ PAT ↓ |
| **A×P** | MAT ↓ | 19.6 | 22.7 | 1.4 | 4.7 | 0.4 | 0.9 | 0.0 | 0.0 |
|  | MAT ↑ | 19.8 | 22.8 | 1.4 | 4.7 | 0.4 | 0.9 | 0.0 | 0.0 |
|  | PAT ↓ | 1.5 | 1.7 | 0.1 | 0.4 | 0.0 | 0.1 | 0.0 | 0.0 |
|  | PAT ↑ | 4.2 | 4.8 | 0.3 | 1.0 | 0.1 | 0.2 | 0.0 | 0.0 |
|  | MATCH ↓↓ | 0.6 | 0.7 | 0.0 | 0.1 | 0.0 | 0.0 | 0.0 | 0.0 |
|  | MATCH ↑↑ | 0.9 | 1.0 | 0.1 | 0.2 | 0.0 | 0.0 | 0.0 | 0.0 |
|  | MAT ↓ PAT ↑ | 0.1 | 0.1 | 0.0 | 0.0 | 0.0 | 0.0 | 0.0 | 0.0 |
|  | MAT ↑ PAT ↓ | 0.2 | 0.2 | 0.0 | 0.0 | 0.0 | 0.0 | 0.0 | 0.0 |

**Table S6.** Empirical contingency table of shared DPE genes between the reciprocal crosses C×P and P×C.

|  |  | P×C |  |  |  |  |  |  |  |
| --- | --- | --- | --- | --- | --- | --- | --- | --- | --- |
|  | DPE category | MAT ↓ | MAT ↑ | PAT ↓ | PAT ↑ | MATCH ↓↓ | MATCH ↑↑ | MAT ↓ PAT ↑ | MAT ↑ PAT ↓ |
| C×P | MAT ↓ | 11 | 2 | **21** | 6 | 6 |  |  |  |
|  | MAT ↑ | 11 | 4 | 12 | **27** | 4 |  | 2 |  |
|  | PAT ↓ | **3** | 0 | 1 | 2 |  |  |  |  |
|  | PAT ↑ | 1 | **6** | 6 | 3 |  | 1 |  |  |
|  | MATCH ↓↓ |  |  |  | 1 | 1 |  |  |  |
|  | MATCH ↑↑ |  |  | 2 |  |  | 2 |  |  |
|  | MAT ↓ PAT ↑ |  |  |  |  |  |  |  |  |
|  | MAT ↑ PAT ↓ |  |  |  |  |  |  |  |  |

**Table S7.** ***Expected numbers* of shared DPE genes between the reciprocal crosses C×P and P×C, given the observed numbers in Table 1**.

|  |  | P×C |  |  |  |  |  |  |  |
| --- | --- | --- | --- | --- | --- | --- | --- | --- | --- |
|  | DPE category | MAT ↓ | MAT ↑ | PAT ↓ | PAT ↑ | MATCH ↓↓ | MATCH ↑↑ | MAT ↓ PAT ↑ | MAT ↑ PAT ↓ |
| **C×P** | MAT ↓ | 9.5 | 7.0 | 14.9 | 15.3 | 3.5 | 2.6 | 0.3 | 0.3 |
|  | MAT ↑ | 11.0 | 8.1 | 17.2 | 17.7 | 4.1 | 3.0 | 0.4 | 0.3 |
|  | PAT ↓ | 0.7 | 0.5 | 1.1 | 1.1 | 0.3 | 0.2 | 0.0 | 0.0 |
|  | PAT ↑ | 2.3 | 1.7 | 3.5 | 3.6 | 0.8 | 0.6 | 0.1 | 0.1 |
|  | MATCH ↓↓ | 0.2 | 0.1 | 0.3 | 0.3 | 0.1 | 0.0 | 0.0 | 0.0 |
|  | MATCH ↑↑ | 0.4 | 0.3 | 0.7 | 0.7 | 0.2 | 0.1 | 0.0 | 0.0 |
|  | MAT ↓ PAT ↑ | 0.0 | 0.0 | 0.0 | 0.0 | 0.0 | 0.0 | 0.0 | 0.0 |
|  | MAT ↑ PAT ↓ | 0.0 | 0.0 | 0.0 | 0.0 | 0.0 | 0.0 | 0.0 | 0.0 |

**Table S8.** Empirical contingency table of shared DPE genes between the reciprocal crosses A×P and P×A.

|  |  | P×A |  |  |  |  |  |  |  |
| --- | --- | --- | --- | --- | --- | --- | --- | --- | --- |
| A×P | DPE category | MAT ↓ | MAT ↑ | PAT ↓ | PAT ↑ | MATCH ↓↓ | MATCH ↑↑ | MAT ↓ PAT ↑ | MAT ↑ PAT ↓ |
|  | MAT ↓ | **25** | 2 | **44** | 17 | 15 | 2 |  | 1 |
|  | MAT ↑ | 15 | **13** | 24 | **73** | 4 | 2 | 1 |  |
|  | PAT ↓ | 1 |  | 2 | **6** | 2 | 0 |  |  |
|  | PAT ↑ | 1 | 6 | **15** | 2 | 3 | 1 |  | 1 |
|  | MATCH ↓↓ | 2 | 0 | 1 | 1 | 2 |  |  |  |
|  | MATCH ↑↑ | 1 | 3 | 2 | 1 | 1 |  |  |  |
|  | MAT ↓ PAT ↑ |  | 1 |  |  |  |  |  |  |
|  | MAT ↑ PAT ↓ |  |  | 1 | 1 |  |  |  |  |

**Table S9.**
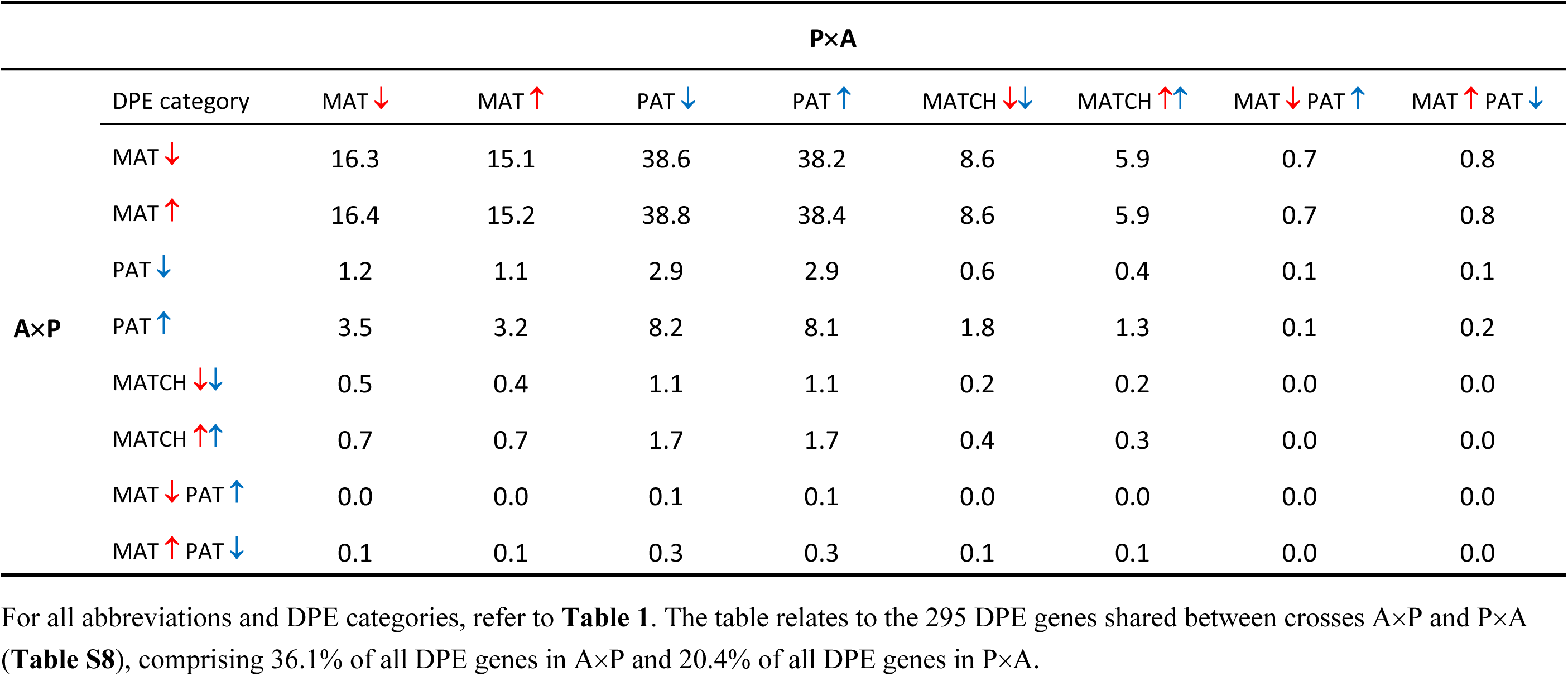
***Expected numbers* of shared DPE genes between the reciprocal crosses A×P and P×A, given the observed numbers in Table 1**.

**Table S10.** Empirical contingency table of shared DPE genes between the reciprocal crosses A×C and C×A.

|  |  | C×A |  |  |  |  |  |  |  |
| --- | --- | --- | --- | --- | --- | --- | --- | --- | --- |
|  | DPE category | MAT ↓ | MAT ↑ | PAT ↓ | PAT ↑ | MATCH ↓↓ | MATCH ↑↑ | MAT ↓ PAT ↑ | MAT ↑ PAT ↓ |
| A×C | MAT ↓ | 4 | 7 | 9 | 1 | 1 |  | 1 |  |
|  | MAT ↑ | 7 | 4 |  | **33** |  | 1 | 2 |  |
|  | PAT ↓ | 5 | 0 | 3 | 4 |  |  |  |  |
|  | PAT ↑ | 1 | **32** | 4 | 2 |  |  |  | 1 |
|  | MATCH ↓↓ | 3 | 0 | 1 |  |  |  |  |  |
|  | MATCH ↑↑ | 0 | 2 |  |  |  | 1 |  |  |
|  | MAT ↓ PAT ↑ |  | 2 | 1 |  | 1 |  |  |  |
|  | MAT ↑ PAT ↓ |  |  |  | 1 |  |  |  |  |

**Table S11.** ***Expected numbers* of shared DPE genes between the reciprocal crosses A×C and C×A, given the observed numbers in Table 1**.

|  |  | C×A |  |  |  |  |  |  |  |
| --- | --- | --- | --- | --- | --- | --- | --- | --- | --- |
|  | DPE category | MAT ↓ | MAT ↑ | PAT ↓ | PAT ↑ | MATCH ↓↓ | MATCH ↑↑ | MAT ↓ PAT ↑ | MAT ↑ PAT ↓ |
| **A×C** | MAT ↓ | 6.3 | 13.4 | 8.9 | 11.3 | 0.6 | 0.8 | 0.3 | 0.2 |
|  | MAT ↑ | 8.2 | 17.6 | 11.6 | 14.8 | 0.8 | 1.0 | 0.4 | 0.3 |
|  | PAT ↓ | 1.9 | 4.0 | 2.7 | 3.4 | 0.2 | 0.2 | 0.1 | 0.1 |
|  | PAT ↑ | 3.0 | 6.5 | 4.3 | 5.5 | 0.3 | 0.4 | 0.1 | 0.1 |
|  | MATCH ↓↓ | 0.2 | 0.5 | 0.3 | 0.4 | 0.0 | 0.0 | 0.0 | 0.0 |
|  | MATCH ↑↑ | 0.3 | 0.6 | 0.4 | 0.5 | 0.0 | 0.0 | 0.0 | 0.0 |
|  | MAT ↓ PAT ↑ | 0.2 | 0.4 | 0.2 | 0.3 | 0.0 | 0.0 | 0.0 | 0.0 |
|  | MAT ↑ PAT ↓ | 0.1 | 0.1 | 0.1 | 0.1 | 0.0 | 0.0 | 0.0 | 0.0 |

**Table S12.**
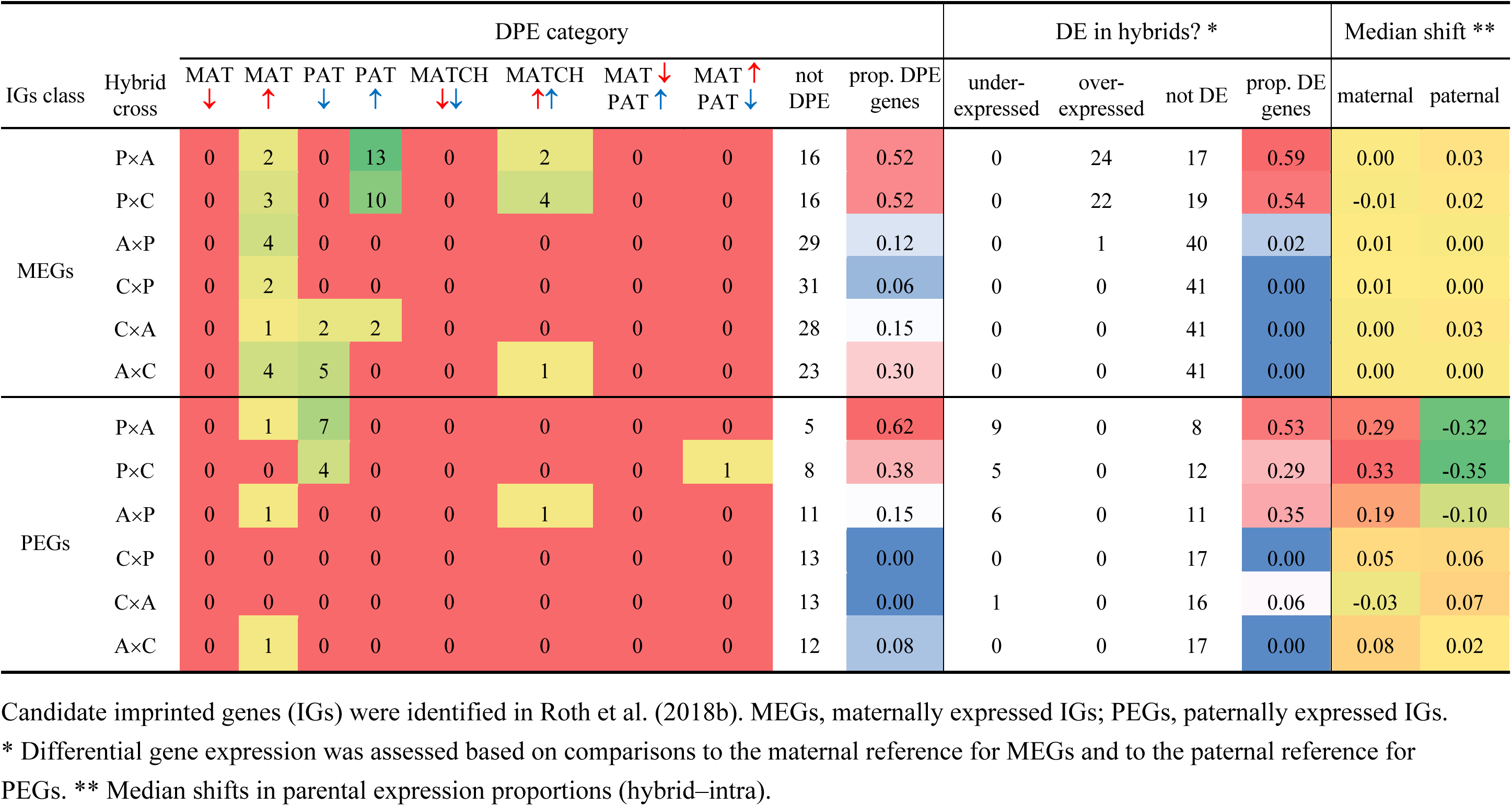
Differential parental (DPE) and total (DE) expression changes among 58 conserved imprinted wild tomato genes.

## Supplementary Figures

**Figure S1.**
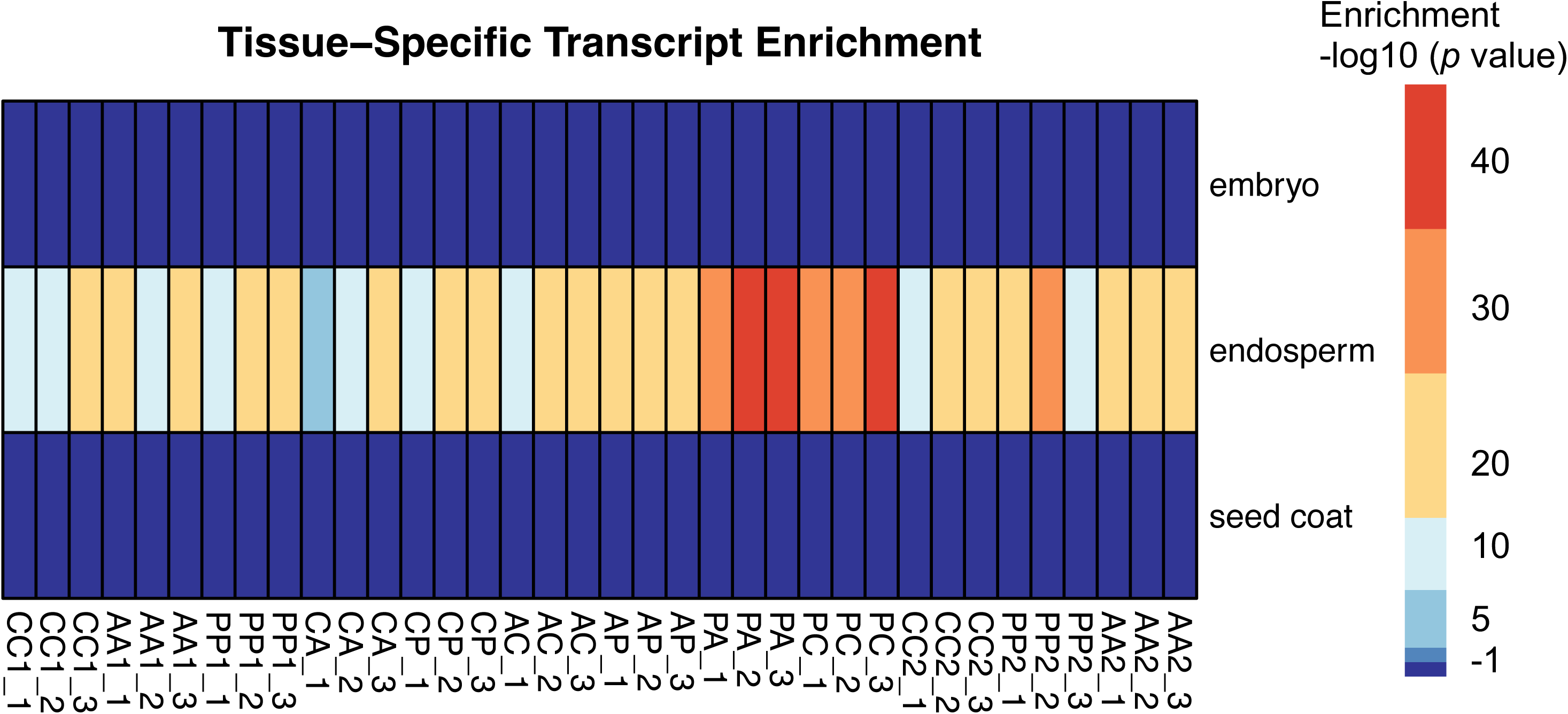
Heat map representing tissue-specific enrichment for each of our 36 endosperm samples (following Schon and Nodine 2017, using data from Pattison et al. 2015). Note the absence of any evidence for possible contamination with embryo or seed coat tissue in our laser-dissected endosperm data.

**Figure S2.**
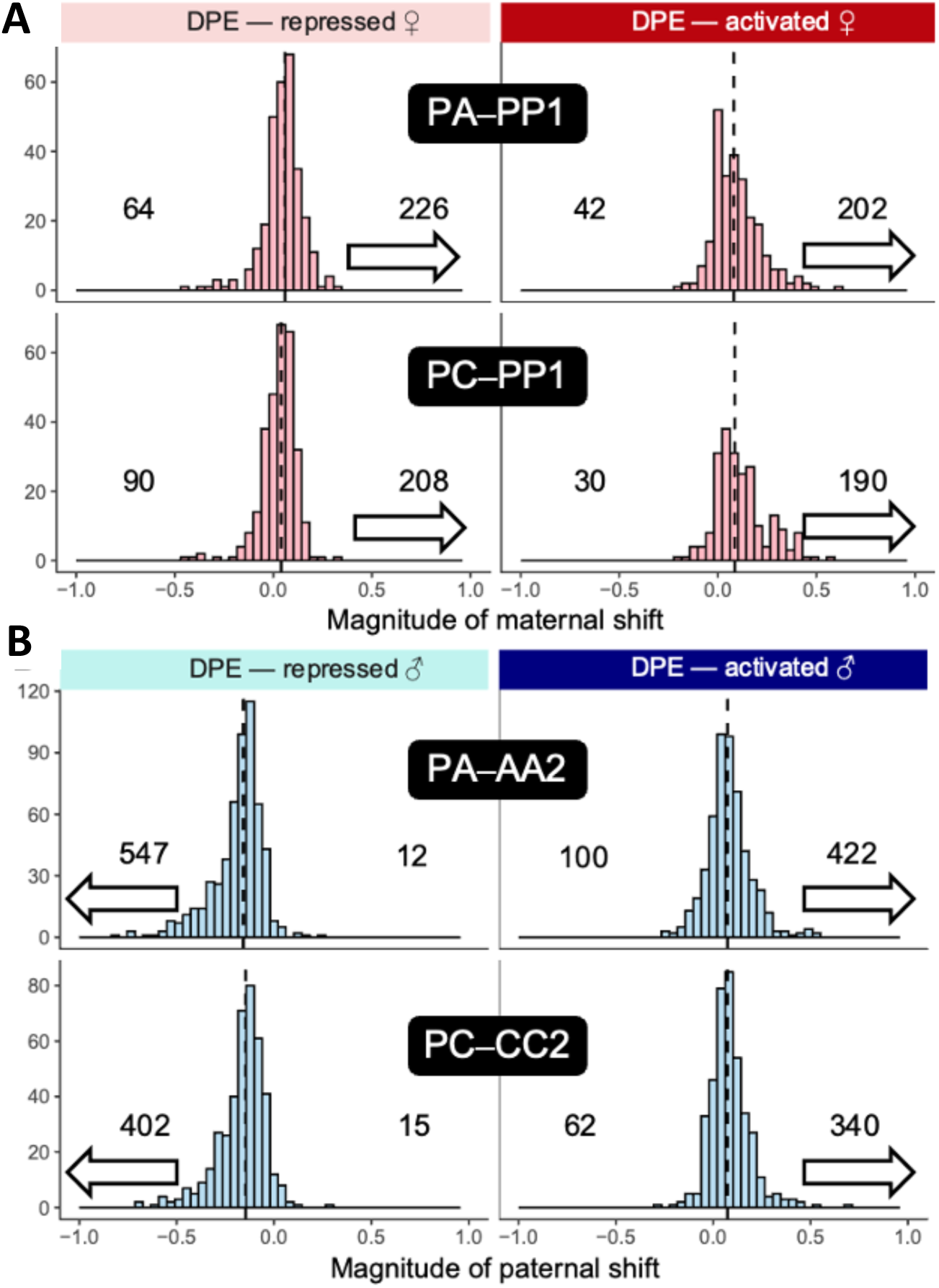
Distribution of parental expression shifts for differentially expressed (DE) genes between hybrid and intraspecific reference data (left-hand panels) and differentially parentally expressed (DPE) genes (right-hand panels) in ME-like crosses PA and PC. **A**, distribution of maternal shift in under-and overexpressed (hybrid–intra) DE genes in PA–PP1 and PC–PP1 comparisons. **B**, distribution of paternal shift in under- and overexpressed (hybrid–intra) DE genes in PA–AA2 and PC–CC2 comparisons. **C**, distribution of maternal shift in maternally repressed and -activated (hybrid–intra) DPE genes in PA–PP1 and PC–PP1 comparisons. **D**, distribution of paternal shift in paternally repressed and -activated (hybrid–intra) DPE genes in PA–AA2 and PC–CC2 comparisons. The dashed vertical lines correspond to the median shift value of each distribution. In **A** and **B**, the numbers refer to genes with shift values more negative than -0.10 (left) and more positive than 0.10 (right), with white arrows indicating the major direction. In **C** and **D**, the numbers refer to all genes with non-zero shift value. Analogous estimates for PE-like and partially viable hybrid crosses are presented in **Fig. S3**.

**Figure S3.**
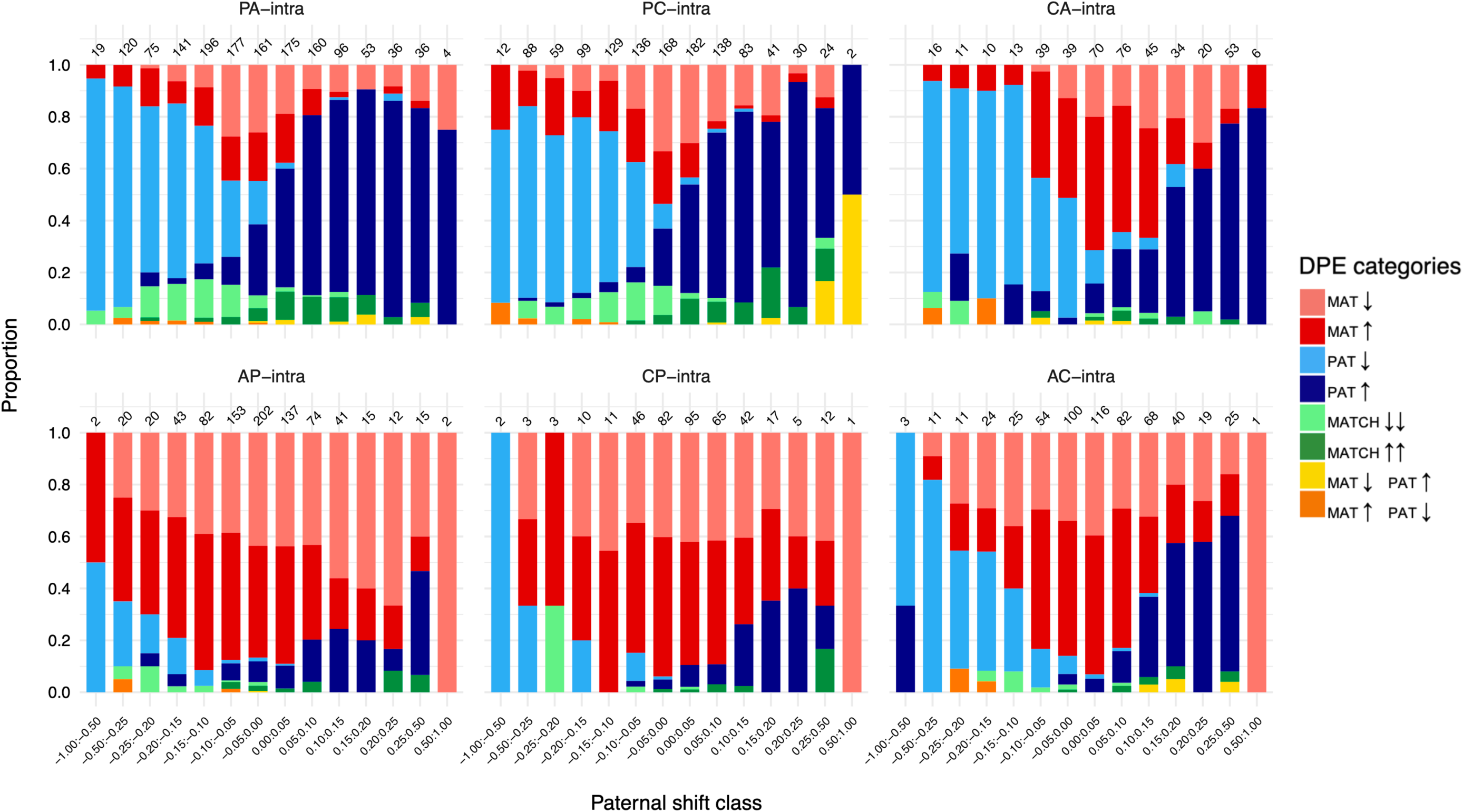
Maternal and paternal expression changes per paternal shift class in the comparison between hybrid endosperms and their corresponding intraspecific reference. Numbers at the top of the bars show the total number of DPE genes in each shift class (bin). A common set of 5,015 genes was assessed in each of the six comparisons and only genes with DPE are shown. Note that paternal shift classes between –0.25 and 0.25 have interval size 0.05 but that outside this range, intervals are larger, as indicated along the bottom x-axis.

